# Distinct epicardial gene regulatory programmes drive development and regeneration of the zebrafish heart

**DOI:** 10.1101/2021.06.29.450229

**Authors:** Michael Weinberger, Filipa C. Simões, Tatjana Sauka-Spengler, Paul R. Riley

## Abstract

Unlike the adult mammalian heart, which has limited regenerative capacity, the zebrafish heart can fully regenerate following injury. Reactivation of cardiac developmental programmes is considered key to successfully regenerating the heart, yet the regulatory elements underlying the response triggered upon injury and during development remain elusive. Organ-wide activation of the epicardium is essential for zebrafish heart regeneration and is considered a potential regenerative source to target in the mammalian heart. Here we compared the transcriptome and epigenome of the developing and regenerating zebrafish epicardium by integrating gene expression profiles with open chromatin ATAC-seq data. By generating gene regulatory networks associated with epicardial development and regeneration, we inferred genetic programmes driving each of these processes, which were largely distinct. We identified wt1a, wt1b, and the AP-1 subunits junbb, fosab and fosb as central regulators of the developing network, whereas hif1ab, zbtb7a, tbx2b and nrf1 featured as putative central regulators of the regenerating epicardial network. By interrogating developmental gene regulatory networks that drive cell-specific transcriptional heterogeneity, we tested novel subpopulation-related epicardial enhancers *in vivo.* Taken together, our work revealed striking differences between the regulatory blueprint deployed during epicardial development and regeneration. These findings challenge the dogma that heart regeneration is essentially a reactivation of developmental programmes, and provide important insights into epicardial regulation that can assist in developing therapeutic approaches to enable tissue regeneration in the adult mammalian heart.

## Introduction

In mammals, the capacity to repair the heart following injury rapidly declines soon after birth (Haubner, et al., 2016; Porrello, et al., 2011). In contrast, the zebrafish heart can regenerate up to and throughout adulthood (Gonzalez-Rosa, et al., 2011; Poss, et al., 2002), offering the possibility to investigate the mechanisms underlying cardiac regeneration (Reviewed in (Gonzalez-Rosa, et al., 2017; Foglia and Poss, 2016)). The epicardium, a mesothelial layer of cells enveloping the vertebrate heart muscle, is essential for heart development (Simoes and Riley, 2018) and critically required for zebrafish heart regeneration (Wang, et al., 2015). While epicardial cells in the homeostatic adult heart are mostly dormant, they become rapidly activated following cardiac injury. Activated epicardial cells re-enter the cell cycle and re-express developmental genes such as the transcription factors *tbx18* and *wt1b*, as well as the retinoic acid synthesizing enzyme *aldh1a2* (Gonzalez-Rosa, et al., 2012; Gonzalez-Rosa, et al., 2011; Kikuchi, et al., 2011a; Kikuchi, et al., 2011b; Lepilina, et al., 2006). Initially, epicardial cell activation is an organ-wide response fully manifested by 3 days post-injury (dpi) and then becomes restricted to the site of injury by 7 dpi (Gonzalez-Rosa, et al., 2011; Lepilina, et al., 2006). Activated epicardial cells are amongst the first to migrate into the injury area, depositing extracellular matrix (ECM) components such as fibronectin and collagens (Marro, et al., 2016; Mercer, et al., 2013; Wang, et al., 2013). This ECM scaffold supports heart regeneration and guides the migration of cardiomyocytes into the wound region (Wang, et al., 2015; Wang, et al., 2013). Epicardium-derived chemokines such as cxcl12a also promote the growth of the myocardium in the regenerating area (Itou, et al., 2012) and the epicardium further stimulates myocardial proliferation following injury (Wang, et al., 2015) by secreting mitogenic factors such as retinoic acid (Kikuchi, et al., 2011b), sonic hedgehog (Sugimoto, et al., 2017; Choi, et al., 2013), insulin-like growth factor 2b (igf2b) (Huang, et al., 2013), bmp2b (Wu, et al., 2016) and tgfβ (Chablais and Jazwinska, 2012). Finally, epicardial cells can also promote heart regeneration directly by undergoing epithelial-to-mesenchymal transition (EMT), invading the subepicardial tissue as epicardium-derived cells (EPDCs) (Simoes and Riley, 2018) and giving rise to perivascular cells and fibroblasts (Gonzalez-Rosa, et al., 2012). Epicardial EMT is regulated by fibroblast growth factors (Lepilina, et al., 2006) and platelet-derived growth factors (Kim, et al., 2010), whereas hippo signalling regulates the formation of fibroblasts from EPDCs (Xiao, et al., 2018). Thus, the epicardium supports heart regeneration both directly and indirectly, and a growing number of signalling pathways have been implicated in mediating the epicardial regulation of other cardiac tissues.

Across species, recapitulation of developmental programmes has generally been considered a key feature to successfully regenerate the heart following injury (Goldman and Poss, 2020; Wang, et al., 2020b; Wang, et al., 2019). Moreover, a developmental response to injury or pathology is a more general paradigm, such that even in the diseased adult human heart there is a hallmark of re-expression of embryonic genes, including fetal *α*- and *β*-myosin heavy chain (*α*-MHC and *β*-MHC), brain natriuretic peptide (BNP, *Nppb*) and atrial natriuretic factor (ANF, *Nppa*), associated with pathological remodelling (Dirkx, et al., 2013). In the epicardium, many of the factors expressed by activated adult epicardial cells are also expressed by the developing epicardium, and epicardial functions during heart regeneration and development are very similar (Reviewed in (Simoes and Riley, 2018). However, it has also previously been demonstrated that expression profiles of selected cell-surface markers in mouse embryonic versus injury-activated epicardial cells differ quite significantly (Bollini, et al., 2014). Given the essential roles of the epicardium during both development and regeneration, surprisingly little is known about the upstream transcriptional regulation driving epicardial gene expression programmes during either process. Efficient and precise gene expression is controlled by transcription factors (TFs), which bind to specific DNA sequences within promoters and other distal *cis*-regulatory elements. Enhancers make up the most abundant subset of non-promoter *cis*-regulatory elements: these are short (<1kb) regions of open chromatin, composed of clusters of TF recognition motifs, with the ability to drive transcription over long distances through interaction with cognate promoters (Reviewed (Long, et al., 2016)). Often, multiple enhancers act in a combinatorial fashion to regulate the transcriptional output of a single gene locus. Thus, spatiotemporal tuning of gene expression is modulated by TF presence and enhancer accessibility (Nord, et al., 2013). The activity of genomic regions is dependent on how closely they are structurally associated with the neighbouring nucleosomes, as this affects the capacity of TFs to bind to the DNA sequence (Reviewed in (Felsenfeld and Groudine, 2003)). A weak interaction between a DNA sequence and the surrounding nucleosomes can thus be an indication of TF binding within the sequence. By mapping such open chromatin regions across the genome, it is possible to identify potential active enhancer elements. Assay for Transposase-Accessible Chromatin (ATAC)-sequencing builds on this principle and uses Tn5 transposase to integrate adapter sequences into open chromatin regions, enabling amplification of accessible sequences for next-generation sequencing (Buenrostro, et al., 2013). Recently, regions in the zebrafish genome have been identified as tissue regeneration enhancer elements (TREEs), becoming activated in response to cardiac injury (Begeman, et al., 2020; Wang, et al., 2020a; Goldman, et al., 2017; Pfefferli and Jazwinska, 2017; Kang, et al., 2016). While some epicardial TREEs have been identified in the mouse (Vieira, et al., 2017; Huang, et al., 2012), TREEs of the adult regenerating epicardium have not yet been studied.

Here we compared the transcriptome and epigenome of the developing and regenerating zebrafish epicardium by integrating gene expression and chromatin accessibility ATAC-seq profiles. We identified *bona fide* transcriptional regulators that drive epicardial heterogeneity during cardiac development and showed that putative regulatory regions associated with previously identified epicardial subpopulations (Weinberger, et al., 2020) could drive *in vivo* gene expression in a cell-type-specific manner. We tested the activity of novel subpopulation-specific epicardial enhancers *in vivo* and were able to build three distinct gene regulatory networks that drive epicardial transcriptional heterogeneity during development. We found that the epicardial transcriptomic programmes deployed during development and regeneration are largely distinct, in contrast to current dogma that regeneration largely recapitulates development. By data analysis integration, we have built gene regulatory networks that specifically govern each of these processes. We inferred wt1a, wt1b, and the AP-1 subunits junbb, fosab and fosb as central regulators of the developing epicardial gene regulatory network, while TFs such as hif1ab, zbtb7a, tbx2b and nrf1 featured as putative central regulators of the regenerating epicardial network. Interestingly, even the underlying genetic programmes driving expression of the small cohort of transcripts shared by the developing and the post-injury epicardium were distinct. Further analysis suggested that genomic regulation in the regenerating epicardium allows rapid gene expression onset upon injury, contrasting with the slower genetic blueprint deployed during development.

## Results

### Gene expression profiles of the developing and regenerating zebrafish epicardium are largely distinct

To compare the epicardial gene expression profiles during zebrafish development and following adult heart injury, we analysed global transcriptional landscapes of epicardial cells derived from *TgBAC(tcf21:H2B-Dendra2)^ox182^* and *TgBAC(tcf21:BirA-2a-mCherry*;*βactin:Avi-Cerulean-RanGap)^ox144^* transgenic lines. We obtained bulk RNA-sequencing data from cardiac *tcf21*-positive cells (*tcf21*^+^) and corresponding tcf21-negative (*tcf21*^-^) control cardiac cells at 5 days post fertilisation (dpf), a time point when the epicardium is fully formed. For adult samples, we profiled cardiac *tcf21*^+^ nuclei at 3 days post cryoinjury (dpi) and corresponding sham-control samples (3 dps), using the Biotagging approach to isolate specific nuclei (Simoes, et al., 2020; Trinh, et al., 2017), at a time point when the injured epicardium is highly proliferative (Gonzalez-Rosa, et al., 2011) (Figure 1A). Principal component (PC) analysis revealed that the variability of gene expression between replicates of each condition was low (Figure 1B). The majority of variability was due to differences between larval and adult samples (PC1, 69%), a smaller amount due to differences between larval epicardial and control samples (PC2, 20%). In contrast, adult cryoinjury and sham-control samples were very similar in their gene expression. Directly comparing larval epicardium and adult epicardium following cryoinjury, we found that 8554 genes were significantly enriched in the larval epicardium, while 2472 genes were enriched in the adult cryoinjured epicardium (Figure 1C). Among the genes enriched in the larval epicardium were the chemokine ligand *cxcl14* and the AP-1 transcription factor subunits *fosab*, *junbb* and *jdp2b* (*jun dimerization protein 2b*). The adult cryoinjured epicardium featured several enriched collagens, such as *col1a1a*, *col1a2* and *col12a1b*.

**Figure 1.**
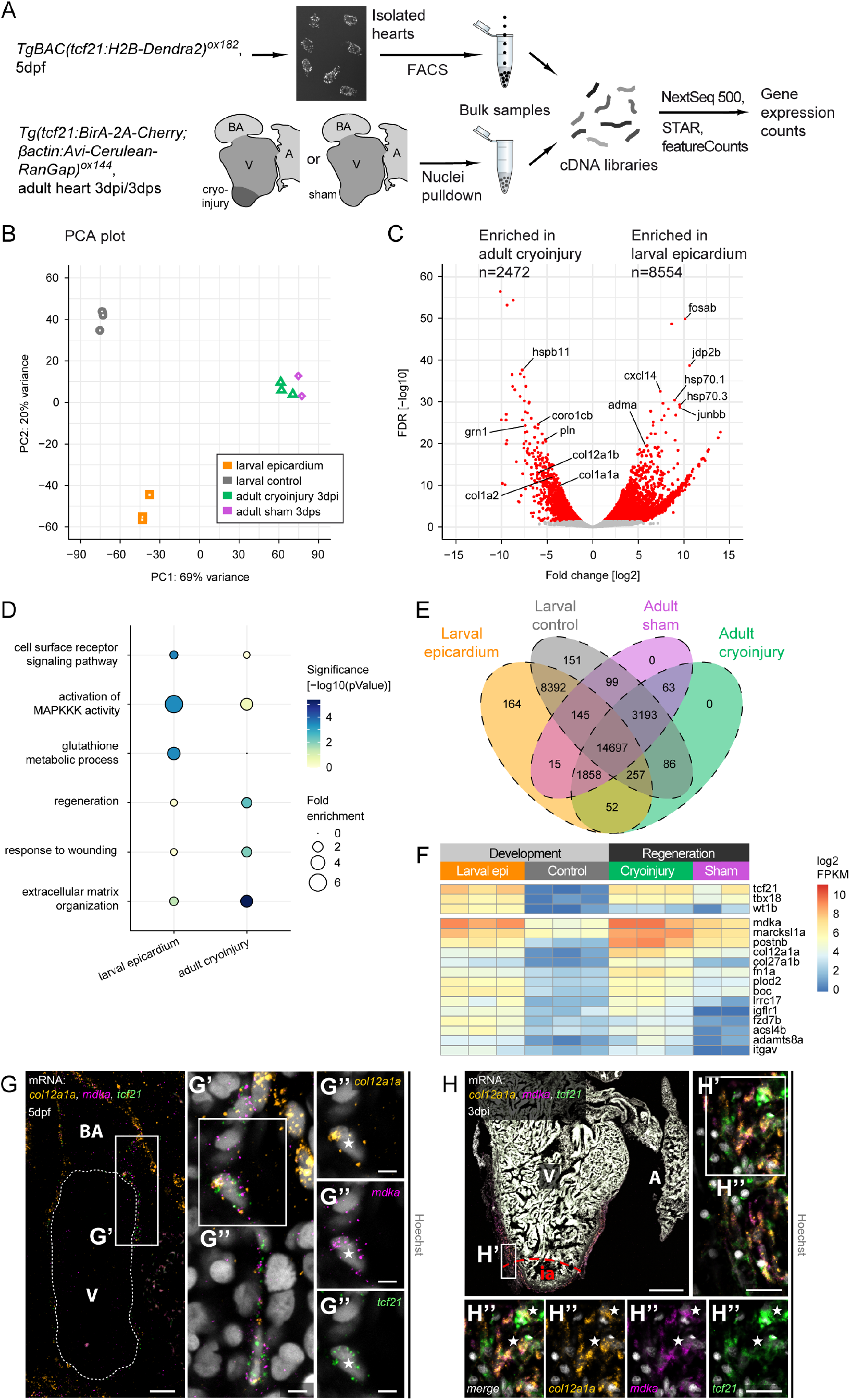
Gene expression programmes in the developing and regenerating zebrafish epicardium are distinct. (A) Overview of the RNA sequencing workflow. Number of biological replicates analysed: larval epicardium n=3, larval control n=3, adult cryo n=3, adult sham n=2. (B) Principal component clustering of transcriptome samples. (C) Differential gene expression analysis of larval epicardium versus adult cryoinjured epicardium. Shown are log2 transformed expression fold changes and Benjamini-Hochberg adjusted Wald test p-values for each gene. Significantly enriched genes (adjusted p-value < 0.05) are coloured in red. Indicated are the numbers of enriched genes in each condition. (D) GO term over-representation of genes enriched in the larval epicardium (versus adult cryo, left column) and genes enriched in adult cryoinjured epicardium (right column). Bubble size depicts the magnitude of statistical enrichment, colour the significance. (E) Venn diagram depicting the overlap of gene expression enrichment in larval epicardium, larval control, adult cryoinjured and sham-treated epicardium. Indicated are the numbers of genes contained in each intersection. (F) Expression of the epicardial marker genes *tcf21*, *tbx18* and *wt1b*, as well as expression of genes enriched in the larval epicardium over the larval control and enriched in the adult cryoinjured epicardium over the adult sham-injured epicardium. Shown are log2 transformed FPKM values. (G) mRNA staining of *col12a1a* (orange), *mdka* (magenta) and *tcf21* (green) in a 5dpf heart. (G‘,G‘‘) A nucleus (asterisk) in the epicardial region surrounded by *col12a1a*, *mdka* and *tcf21* transcripts. (H) mRNA staining of *col12a1a* (orange), *mdka* (magenta) and *tcf21* (green) in a 3dpi cryoinjured heart. ia=injury area. (H‘,H‘‘) Single nuclei (asterisks) in the epicardial region surrounded by *col12a1a*, *mdka* and *tcf21* transcripts. Scale bars H: 100µm, G,H’,H’’: 20µm, G’,G’’: 5µm. Colour channels adjusted separately for brightness/contrast. G,H are single optical sections. dpf=days-post-fertilisation, dpi=days-post-injury, V=ventricle, A=atrium, BA=bulbus arteriosus. See also Fig. S1.

Genes enriched in the developing epicardium were associated with multiple gene ontology (GO) terms related to cellular signalling, such as “cell surface receptor signalling pathway” (p=0.00029) and “activation of MAPKKK activity” (p=0.00037) (Figure 1D). Genes enriched in the regenerating epicardium were mainly associated with injury-related processes such as “regeneration” (p=0.00384), “response to wounding” (p=0.00972) and extracellular matrix organization (p=4.2*10-6).

After assessing the transcriptomic differences between the developing and the regenerating epicardium, we next asked whether there were common or distinct gene programmes that characterised each of these processes. To this end, we compared gene expression in the larval epicardium to that in the non-epicardial larval control cells and intersected this comparison with that of adult cryoinjured versus sham-control epicardium (Figure 1E). The number of genes expressed in the larval epicardium (n=28877) was approximately 44% higher than in the regenerating cryoinjured epicardium (n=20012), indicating higher transcriptomic diversity of the larval epicardium. Most genes expressed in the adult epicardial samples were not differentially expressed between 3 dpi cryoinjured condition and sham-control. This common early transcriptomic response is supported by sample similarity shown by our PC analysis (Figure 1B) and is most probably due to the general downstream effects of cutting open the pericardial cavity to expose the ventricular chamber in both sham and injury-inducing procedures (Gonzalez-Rosa, et al., 2011). Yet, a cohort of 52 genes enriched in the adult cryoinjured versus sham-control epicardium was also expressed and enriched in the larval epicardium versus the non-epicardial larval control (Figure 1E and Table S1). A large proportion of these genes were components of the extracellular matrix and signalling factors (Figure 1F). Using multiplexed hybridization chain reaction (HCR) *in situ* staining (Choi, et al., 2018), we validated the expression of some of the shared transcripts in the developing (Figures 1G,S1A) and cryoinjured epicardium (Figures 1H,S1B). We found *col12a1a* and *midkine a* (*mdka*) genes (Figures 1G,H), as well as *periostin b* (*postnb*) and *plod2* (Figures S1A,S1B) were expressed within the epicardial cell layer, co-localising with *tcf21* transcripts.

In summary, our data suggest that the epicardial transcriptomic response deployed upon adult heart injury at 3 dpi is not a mere recapitulation of the epicardial gene programme acting during development, as a large number of genes were differentially enriched in each setting (see Figure S1C for top genes enriched specifically in either larval or cryoinjured epicardium). However, we also identified several transcripts shared by both the developing and the regenerating epicardium, suggesting that these genes might represent essential regulators of epicardial function in both settings.

### Chromatin accessibility profiles of the developing and regenerating epicardium

Next, we sought to gain insight into the genomic regulation that underlies epicardial gene expression during heart development and regeneration. To do so, we investigated chromatin accessibility by performing Assay for Transposase-Accessible Chromatin with high-throughput sequencing (ATAC-seq) on FAC-sorted 5 dpf *tcf21*^+^ epicardial cells (isolated from *TgBAC(tcf21:H2B-Dendra2)^ox182^* transgenic line) and corresponding *tcf21*^-^ cardiac controls cells, 5 dpf *tbx18*^+^ epicardial cells (isolated from *TgBAC(tbx18:myr-Citrine)^ox185^* transgenic line) and corresponding *tbx18*^-^ cardiac control cells, and 3 dpi regenerating *tcf21*^+^ epicardial cells and *tcf21*^+^ cells from corresponding sham-control samples (isolated from *TgBAC(tcf21:BirA-2a-mCherry)^ox143^* transgenic line) (Figure 2A).

**Figure 2.**
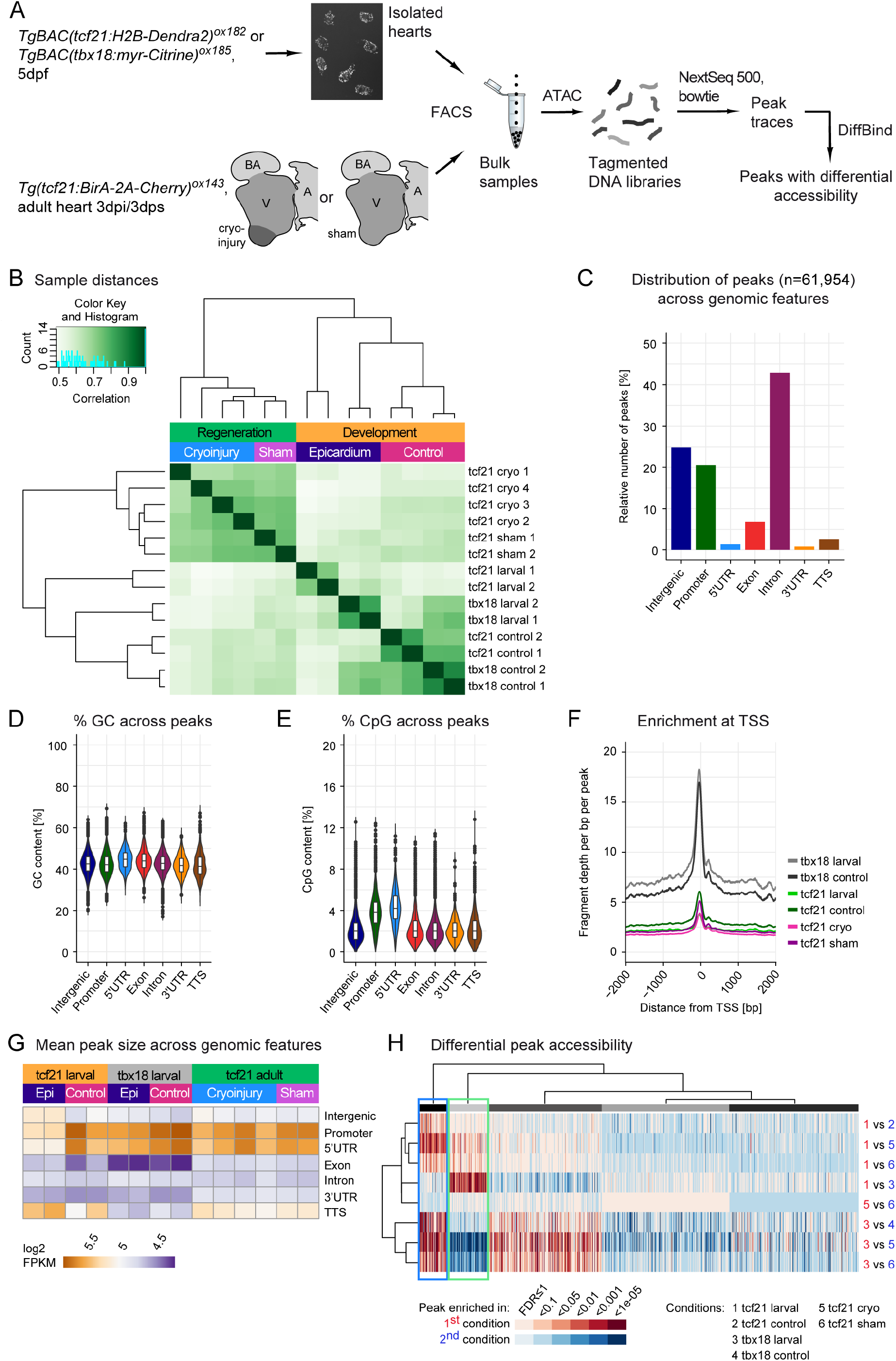
Chromatin accessibility in the developing and regenerating zebrafish epicardium. (A) Overview of the ATAC sequencing workflow. (B) Distance-based clustering of the Pearson correlations of larval and adult accessible chromatin region profiles, as indicated by the dendrogram. Correlation values are indicated by colour. (C) Relative quantification of accessible chromatin region (peak) distribution across genomic features. (D) Relative quantification of peak GC content across genomic features. (E) Relative quantification of peak CpG content across genomic features. (F) Average relative sequencing read densities at transcriptional start sites. Distance from the TSS is shown in base pairs (bp). (G) Average peak size, categorised according to genomic feature overlap. Shown are log2 transformed FPKM values. (H) Overview of differential peak accessibility. Columns show peaks, rows show pairwise comparisons between conditions, as indicated by the numbers on the right hand side of the heatmap and in the legend at the bottom. Colours indicate adjusted p-values, red colours show enrichment in the first condition of a comparison, blue colours show enrichment in the second condition. Peaks are k-means clustered, as indicated by grey colour bars. K-means clusters are hierarchically clustered on the row-wise average values within each cluster. Peaks enriched in tcf21 larval and tbx18 larval are highlighted by a blue box, peaks enriched in tcf21 larval and tcf21 cryo by a green box. dpf=days-post-fertilisation, dpi=days-post-injury, V=ventricle, A=atrium, BA=bulbus arteriosus. Number of biological replicates analysed: larval epicardium n=2, larval control n=2, adult cryo n=4, adult sham n=2. Box plots in D,E represent median, first and third quartiles.

Following identification of accessible chromatin regions (peaks) across the genome via MACS2 (Zhang, et al., 2008), we clustered all samples according to the similarity of their peak positions via DiffBind (Stark and Brown, 2011) (Figure 2B). The biological replicates of the same condition clustered closely together. All *tcf21*^+^ epicardial samples clustered together, with samples from cryoinjured hearts (tcf21 cryo) being very similar to those of sham-control hearts (tcf21 sham), similar to the transcriptomic datasets (Figure 1B). While the peak profiles of *tcf21*^+^ larval epicardial samples (tcf21 larval) shared little similarity with those of *tcf21*^-^ larval cardiac control cells (tcf21 control), peak profiles of *tbx18*^+^ larval epicardial samples (tbx18 larval) showed a higher correlation with those of both *tcf21*^-^ and *tbx18*^-^ larval control conditions. This aligns with our previous observation that the *tbx18*^+^, *tcf21*^-^ Epi2 cell population forms part of the developing bulbus arteriosus (BA) smooth muscle layer, alongside *tbx18*^-^ and *tcf21*^-^ cells (Weinberger, et al., 2020).

We next constructed a consensus peak set of 61,954 accessible regions across samples of all conditions (Stark and Brown, 2011) and analysed peak genomic features using Homer (Heinz, et al., 2010). 20% of the peaks resided within promoters, 43% in introns, 7% overlapped annotated exons, and 25% of the peaks were located in intergenic regions (Figure 2C). The median GC content varied between 42% and 44% across peaks overlapping different genomic features (Figure 2D). The median peak sequence proportion occupied by elongated GC-rich regions (CpG islands) ranged from 2% to 4%, with peaks in promoters and 5’ untranslated regions (5’UTRs) displaying the highest median CpG content (Figure 2E). Furthermore, we analysed the number of sequencing reads aligning to regions surrounding transcriptional start sites (TSS) in the genome (Figure 2F). In all conditions, the average read coverage across replicates was elevated at TSSs, with particular enrichment in the *tbx18*^+^ larval and *tbx18^-^* control conditions. We computed average peak sizes across conditions and genomic features and ascertained that peaks located within promoters or directly downstream of them were generally larger when compared to peaks overlapping other genomic features (corrected p-values t-test promoter versus non-promoter peaks: tcf21 larval 1 5.5*10^-15^, tcf21 larval 2 4.5*10^-11^, tcf21 control 1 <2.2*10^-16^, tcf21 control 2 1.2*10^-124^, tbx18 larval 1 9*10^-182^, tbx18 larval 2 1.9*10^-227^, tbx18 control 1 3.1*10^-299^, tbx18 control 2 <2.2*10^-16^, tcf21 cryo 1 2.3*10^-119^, tcf21 cryo 2 1.2 *10^-294^, tcf21 cryo 3 <2.2*10^-16^, tcf21 cryo 4 6.7*10^-293^, tcf21 sham 1 <2.2*10^-16^, tcf21 sham 2 <2.2*10^-16^) (Figure 2G). However, in the *tcf21*^+^ larval epicardium peaks located in the proximity of the transcriptional termination site (TTS) possessed, on average, the highest accessibility. Furthermore, the average intergenic peak size was larger in the *tcf21*^+^ larval epicardium than in the other conditions. These observations suggested that the DNA regulatory elements underlying gene expression programmes of the developing epicardium and the injured adult epicardium were differentially accessible.

To investigate the regulatory differences between conditions on a peak-by-peak basis, we performed pairwise differential peak accessibility (DPA) analyses using DESeq2 (Love, et al., 2014). By clustering peaks according to their DPA profiles, we identified a group that showed enriched accessibility in both tcf21 and tbx18 larval conditions, compared to tcf21 cryo samples (Figure 2H, blue box). In contrast, a different group of peaks was enriched in both tcf21 larval and tcf21 cryo conditions when compared to tbx18 larval samples (Figure 2H, green box). The first group of peaks thus appears to play a role across the larval epicardium, but not in the adult regenerating counterpart, whereas the latter group might be important in the *tcf21*^+^ epicardium, but not in the developing *tbx18*^+^ epicardium.

### Differential gene regulation underlies transcriptomic heterogeneity in the developing zebrafish epicardium

We first focused our ATAC-seq data analysis on the direct comparison between the *tcf21*^+^ and the *tbx18*^+^ larval epicardium to uncover the regulatory regions governing transcriptomic heterogeneity in the developing epicardium. We identified 5834 peaks (9% of total peak set) significantly enriched (adjusted p-value < 0.05) in epicardial cells expressing *tcf21* and 2617 peaks (4%) enriched in epicardial cells expressing *tbx18* (Figure 3A). Annotating peaks to the closest expressed gene in the respective condition with Homer (Heinz, et al., 2010), we found that 5 out of 7 peaks annotated to *tcf21* locus were among those enriched in the *tcf21*^+^ epicardium and 3 out of 11 peaks annotated to *tbx18* were enriched in *tbx18*^+^ epicardial cells, suggesting these elements might be involved in the regulation of *tcf21* and *tbx18*, respectively. None of the peaks annotated to *tcf21* were enriched in the *tbx18*^+^ epicardium, and vice versa.

**Figure 3.**
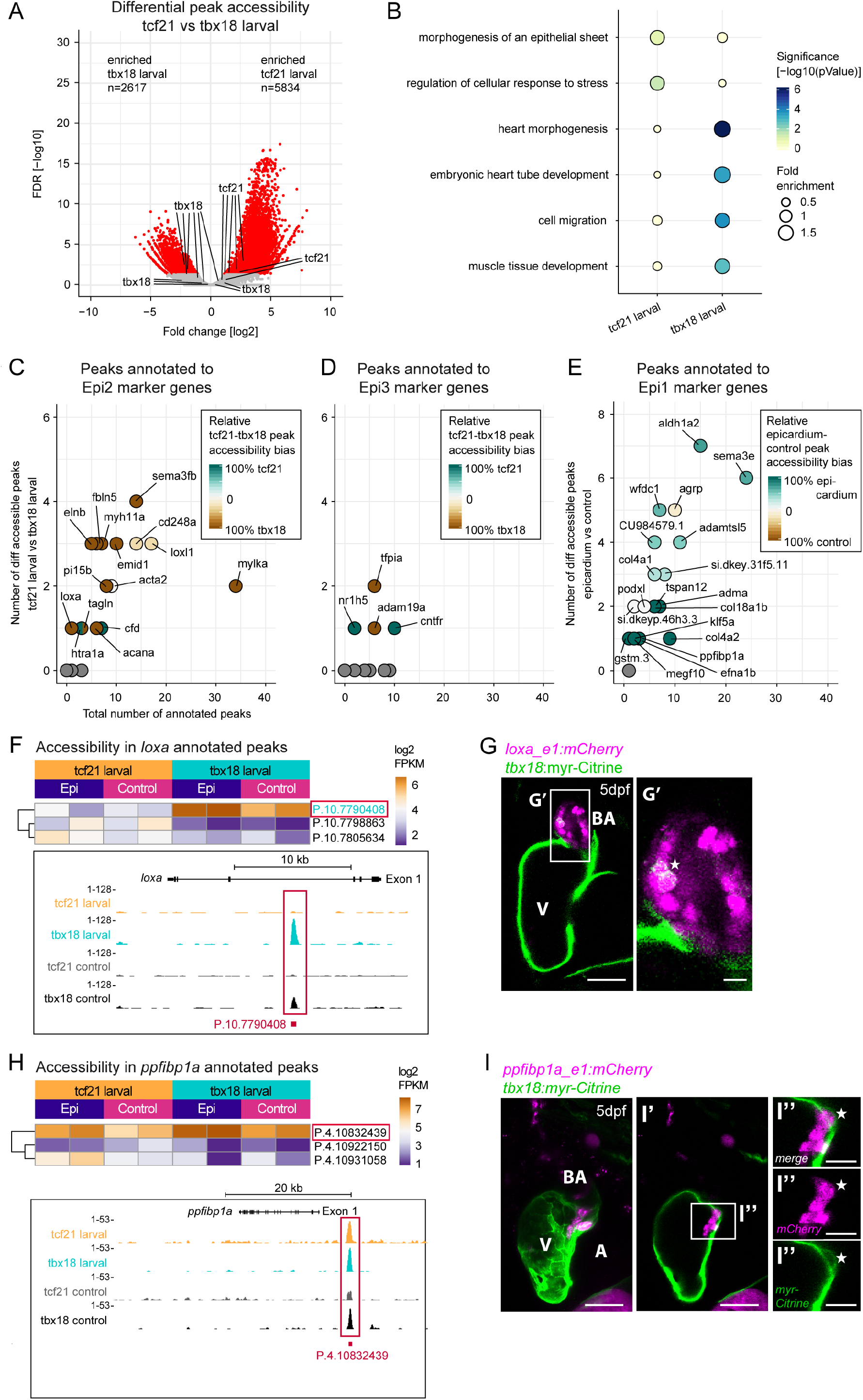
Chromatin accessibility in the developing zebrafish epicardium is linked to transcriptomic heterogeneity. (A) Differential peak accessibility analysis, comparing *tcf21*^+^ larval epicardium to *tbx18*^+^ larval epicardium. Shown are log2 transformed accessibility fold changes and Benjamini-Hochberg adjusted Wald test p-values for each peak. Significantly enriched peaks (adjusted p-value < 0.05) are coloured in red. Highlighted are peaks close to *tcf21* and *tbx18*. (B) GO term over-representation of genes located close to peaks enriched in the *tcf21*^+^ larval epicardium (left column) or peaks enriched in *tbx18*^+^ larval epicardium (right column). Bubble size depicts the magnitude of statistical enrichment, colour the significance. (C) Differential peak accessibility in the vicinity of Epi2 marker genes (n=20). Bubble colour indicates peak accessibility bias, relating the number of peaks enriched in tcf21 larval to the number of peaks enriched in tbx18 larval. Equal numbers of enriched peaks in each condition, for example, would be indicated by white colour (accessibility bias=0). (D) Differential peak accessibility in the vicinity of Epi3 marker genes (n=20). Bubble colour indicates peak accessibility bias of tcf21 larval to tbx18 larval. (E) Differential peak accessibility in the vicinity of Epi1 marker genes (n=20). Bubble colour indicates peak accessibility bias of epicardium to control. (F) Accessibility of peaks close to the Epi2 marker *loxa*. The heatmap shows log2 transformed FPKM values, the genome tracks show read density. Cyan peak label colour indicates significant enrichment in tbx18 larval. A red frame indicates a peak analysed further in G. (G) loxa_e1 driven *in vivo* reporter expression (magenta) at 5dpf. Expression of *tbx18* is indicated by myr-Citrine fluorescence (green membranes). (G‘) Overlap of loxa_e1 activity and myr-Citrine in the BA (asterisk). (H) Accessibility of peaks close to *ppfibp1a*. A red frame indicates a peak analysed further in I. (I) ppfibp1a_e1 driven *in vivo* reporter expression (magenta) at 5dpf. Expression of *tbx18* is indicated by myr-Citrine fluorescence (green membranes). Close proximity of ppfibp1a_e1 activity and myr-Citrine at the boundary of the ventricle (asterisk). (I‘) Single optical section from I. (I’’) A mCherry,myr-Citrine^+^ cell (asterisk) in the epicardial region. Scale bars in G,I,I’: 50µm, scale bar in G’: 10µm, scale bars in I’’: 20µm. V=ventricle, A=atrium, BA=bulbus arteriosus. See also Fig. S2.

Enriched peaks were annotated to specific genes that featured distinct functional associations. While GO terms such as “Morphogenesis of an epithelial sheet” were associated with the tcf21^+^ epicardium, even though non-significantly (p=0.12), tbx18^+^ population showed significant enrichment in “Heart morphogenesis” (p=8.3e-7), “Cell migration” (p=0.00016) and “Muscle tissue development” (p=0.0016) GO terms (Figure 3B). Thus, the gene regulatory programmes governing the developing *tcf21*^+^ and *tbx18*^+^ epicardial subpopulations appeared to be distinct.

These results suggested a connection between the regulatory programmes identified here and the transcriptionally distinct epicardial subpopulations that we previously identified (Weinberger, et al., 2020). Cells in one of these subpopulations, Epi2, do not express *tcf21* but express *tbx18* and several smooth muscle cells associated genes. We analysed the accessibility of peaks annotated to Epi2 genes and found that most of these markers (13 out of 20) featured peaks significantly enriched in the tbx18^+^ larval condition (Figure 3C). At the same time, very few peaks annotated to Epi2 marker genes were enriched in the tcf21^+^ samples. This is in accordance with Epi2 subpopulation expressing *tbx18* but not *tcf21* and thus indicated that the peaks enriched in tbx18 larval might play a role in activating Epi2-specific gene expression. A different epicardial subpopulation, Epi3, is mainly comprised of cells expressing *tcf21* but not *tbx18* (Weinberger, et al., 2020). However, only a low number of peaks annotated to marker genes of Epi3 was differentially accessible (Figure 3D), possibly due to the small fraction of epicardial cells of Epi3 origin. Finally, most cells in the epicardial subpopulation Epi1 express both *tcf21* and *tbx18* (Weinberger, et al., 2020). Therefore, rather than comparing peak accessibility in *tcf21*^+^ and *tbx18*^+^ epicardium, we combined all larval epicardial samples and compared general peak accessibility in the developing epicardium to larval control samples. We found that most peaks located in the vicinity of Epi1-specific genes were significantly enriched in the larval epicardium, suggesting these putative regulatory sequences might play a role in specifically activating the expression of Epi1-related genes (Figure 3E).

We next asked whether peaks annotated to marker genes of the different epicardial subpopulations could activate gene expression in a cell-type-specific manner *in vivo*. To this end, we cloned chosen putative enhancer regions into our Ac/Ds transposase-based reporter system (Chong-Morrison, et al., 2018). One of the putative regulatory sequences enriched in the *tbx18^+^* epicardium, termed *loxa* enhancer 1 (loxa_e1), was located within intron 3 of the *loxa* gene, an Epi2 marker that encodes extracellular matrix crosslinking enzyme *lysyl oxidase* (Figure 3F). We observed loxa_e1-driven reporter mCherry expression in the BA of the developing zebrafish heart, which co-localised with *tbx18^+^* cells (Figure 3G). We next analysed the activity of a different putative regulatory region, elnb_e2, also enriched in the tbx18^+^ cells and localised approximately 20kb upstream of the extracellular matrix gene *elastin b* (*elnb*), an Epi2-specific gene (Figure S2A). The expression of *elnb* gene is exclusive to the BA (Moriyama, et al., 2016). In accordance, elnb_e2-driven mCherry expression was equally restricted to the larval BA, labelling cells that also expressed *tbx18*-driven Citrine (Figure S2B). The phenotype of cells expressing loxa_e1- and elnb_e2-driven mCherry strongly resembled that of Epi2 cells (Weinberger, et al., 2020), further validating the specificity of the identified regulatory sequences. We next tested the activity of peak regions located in the genomic vicinity of Epi1 marker genes. A peak located approximately 5kb upstream of *ppfibp1a* (ppfibp1a_e1, Figure 3H) was accessible in both tcf21 larval and tbx18 larval conditions. When we cloned this genomic sequence into our Ac/Ds-reporter system, we found ppfibp1a_e-driven mCherry cells that also expressed *tbx18*-driven Citrine (Figure 3I). Testing of a different genomic element, also accessible in both epicardial conditions and located in the promoter region of *sema3e* (sema3e_e12, Figure S2C), showed activity in the epicardium and myocardium (Figure S2D).

In summary, our analysis revealed multiple regulatory programmes active in the developing epicardium, which differentially drive the transcriptional heterogeneity associated with the various larval epicardial subpopulations.

### Epicardial gene expression during development and regeneration depend on distinct regulatory programmes

Given that our data points towards distinct epicardial transcriptomic programmes during development and regeneration, we next sought to investigate the nature of the regulatory programmes underlying such differences. By directly comparing chromatin accessibility in the tcf21 larval and tcf21 cryo conditions, we identified 6330 peaks (10%) with enriched accessibility in the *tcf21*^+^ developing epicardium, and 1811 peaks (3%) with enriched accessibility in the *tcf21*^+^ regenerating epicardium (Figure 4A). 2 out of 8 peaks annotated to *tcf21* gene were among those most significantly enriched in the *tcf21*^+^ regenerating epicardium, with both peaks located at a distance of less than 10kb from the *tcf21* transcriptional start site (Figure 4B). Furthermore, all peaks annotated to *tcf21* showed higher overall accessibility in the regenerating epicardium than in the developing epicardium. This suggests that regulation of *tcf21* expression in the regenerating epicardium might utilise a more complex web of different regulatory elements, allowing for a more rapid response mode upon injury than the more gradual transcriptional regulation deployed during development.

**Figure 4.**
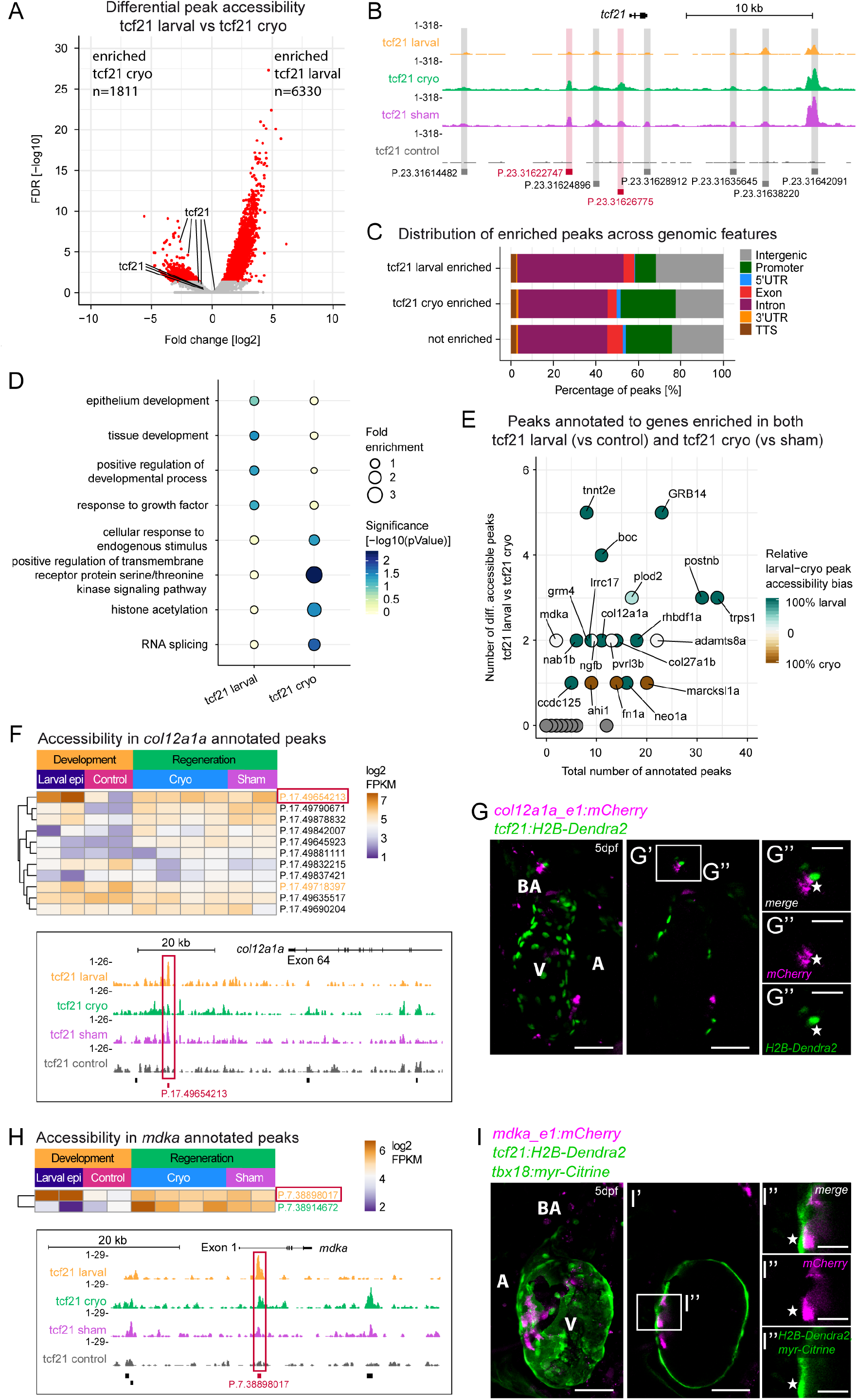
Distinct chromatin accessibility profiles in the developing and regenerating epicardium. (A) Differential peak accessibility analysis, comparing *tcf21*^+^ larval epicardium to *tcf21*^+^ adult cryoinjured epicardium. Shown are log2 transformed accessibility fold changes and Benjamini-Hochberg adjusted Wald test p-values for each peak. Significantly enriched peaks (adjusted p-value < 0.05) are coloured in red. Highlighted are peaks close to *tcf21*. (B) Genome tracks showing chromatin accessibility across the *tcf21* locus. Differentially enriched peaks are highlighted in red, other peaks in grey. (C) Relative quantification of peak distribution across genomic features, categorised according to differential accessibility. (D) GO term over-representation of genes located close to peaks enriched in the *tcf21*^+^ larval epicardium (left column) or peaks enriched in *tcf21*^+^ adult cryoinjured epicardium (right column). Bubble size depicts the magnitude of statistical enrichment, colour the significance. (E) Differential peak accessibility in the vicinity of marker genes of both *tcf21*^+^ larval and adult cryoinjured epicardium (n=34). Bubble colour indicates peak accessibility bias of *tcf21*^+^ larval epicardium to *tcf21*^+^ adult cryoinjured epicardium. (F) Accessibility of peaks close to *col12a1a*. The heatmap shows log2 transformed FPKM values, the genome tracks show read density. A red frame indicates a peak analysed further in G. (G) *col12a1a e1* driven *in vivo* reporter expression (magenta) at 5dpf. Expression of *tcf21* is indicated by H2B-Dendra2 fluorescence (green nuclei). (G‘) Single optical section from G. (G’’) A mCherry^+^ cell (asterisk) in the epicardial region of the BA and in close proximity to a H2B-Dendra2^+^ cell. (H) Accessibility of peaks close to *mdka*. A red frame indicates a peak analysed further in I. (I) *mdka e1* driven *in vivo* reporter expression (magenta) at 5dpf. Expression of *tcf21* is indicated by H2B-Dendra2 fluorescence (green nuclei), expression of *tbx18* by myr-Citrine (green membranes). (I‘) Single optical section from I. (I’’) A mCherry^+^ cell (asterisk) at the boundary of the ventricle and in close proximity to H2B-Dendra2,myr-Citrine^+^ cells. Scale bars in G,G’,I,I’: 50µm, scale bars in G’’,I’’: 20µm. In F and H, yellow peak label colour indicates significant enrichment in tcf21 larval, green colour enrichment in tcf21 cryo. V=ventricle, A=atrium, BA=bulbus arteriosus.

We found that tcf21 larval-enriched peaks at the genomic level featured a lower proportion of promoter peaks than tcf21 cryo enriched peaks (Figure 4C, tcf21 larval: 10%, tcf21 cryo: 26%). In contrast, intronic and intergenic peaks showed the opposite trend (intronic, tcf21 larval: 50%, tcf21 cryo: 42%; intergenic, tcf21 larval: 32%, tcf21 cryo: 23%), suggesting a shift from the distal regulation characterising the developing epicardium towards the proximal, more promoter-based regulation in the regenerating epicardium. We next analysed the functional enrichment of the genes that the differentially accessible genomic regions were associated with. While GO terms such as “tissue development” (p=0.035) and “epithelium development” (p=0.13) were over-represented for tcf21 larval condition (Figure 4D), the top over-represented terms for the tcf21 cryo elements were associated with signalling and gene regulation, such as “positive regulation of transmembrane receptor protein serine/threonine kinase signalling pathway” (p=0.0047) and “histone acetylation” (p=0.033), potentially reflecting an active opening of the epicardial chromatin landscape following injury.

Next, we investigated whether the expression of the 52 transcripts shared by the developing and regenerating epicardium (Table S1) was regulated by similar genomic regions. Notably, 21 of these genes featured peaks significantly enriched in one of the settings, with a strong bias towards enrichment in tcf21 larval samples (Figure 4E). Finally, we tested the activity of peak regions with preferentially enriched accessibility in the larval epicardium but annotated to genes commonly expressed by both the developing and the regenerating epicardium. We found that col12a1a_e1, a region located around 30kb downstream of *col12a1a* (Figure 4F), was active in the developing heart, close to *tcf21*^+^ epicardial cells (Figure 4G). Likewise, a peak region enriched in the larval epicardium and located within the first intron of the *mdka* gene (Figure 4H) displayed activity in the epicardial/subepicardial region of the developing heart at 5dpf (Figure 4I).

Together, our data reveal that gene expression changes observed in the developing and regenerating epicardium are attributable to differences in open chromatin dynamics and highlight unique regulatory regions associated with common transcriptomic programmes of both processes.

### Building developmental-specific and regeneration-specific epicardial gene regulatory networks unravel central regulators of epicardial gene expression

We next sought to understand which transcription factors governed the genomic regulation of epicardial gene expression during heart development and regeneration. For this, we queried our ATAC-seq peak set for the presence of 746 vertebrate transcription factor (TF) binding motifs obtained from the JASPAR 2020 database (Fornes, et al., 2020). We jointly analysed chromatin accessibility and motif occurrence using chromVAR (Schep, et al., 2017) and clustered our ATAC-seq samples according to the accessibility deviations of all motifs (termed “motif accessibility”, Figure S3A). Similar to the peak accessibility-based clustering (Figure 2B), adult tcf21 cryo and sham samples clustered closely together (Figure S3A). Also, tcf21 larval samples clustered together and closer to adult tcf21 samples than to tbx18 larval and larval control samples.

We next queried TF motif occupancy in peaks identified as differentially accessible in the developing versus the regenerating epicardium (Figure 5A). We found the increased presence of Fos and Jun binding motifs, subunits of the TF activator protein-1 (AP-1), in peaks preferentially accessible in the injured epicardium (Figure 5A). Interestingly, the AP-1 complex is required for zebrafish heart regeneration and controls chromatin accessibility in cardiomyocytes (Beisaw, et al., 2020). In contrast, Wt1 and KLF2 binding sites were more frequently detected in genomic regions featuring enriched accessibility in the larval context. Tcf21 and TBX18 motifs were present at equal frequencies in both conditions. This analysis suggests that differential chromatin accessibility between the developing and regenerating epicardium may comprise distinct TFs driving epicardial regulatory activity in the embryo and the adult. Therefore, we next asked whether TFs that potentially bind to sites with enriched accessibility in the developing or the regenerating epicardium were preferentially expressed in either of these contexts (Figure 5B). Indeed we found *wt1a* and *wt1b* expression was enriched in the larval epicardium (FDR(*wt1a*)=0.0016, FDR(*wt1b*)=2*10^-8^), matching the increased binding site accessibility of Wt1 in the developing epicardium as compared to the cryoinjured samples. However, we found that the AP-1 complex components *jun*, *junba*, *junbb*, *fosl1a*, *fosab* and *fosb* were all expressed at significantly higher levels in the developing epicardium than in the regenerating epicardium (FDR(*jun*)=9*10^-12^, FDR(*junba*)=5*10^-19^, FDR(*junbb*)=2*10^-29^, FDR(*fosl1a*)=3*10^-5^, FDR(*fosab*)=1*10^-50^, FDR(*fosb*)=3*10^-12^). These findings contrast with the lower accessibility of AP-1 binding sites in larval-enriched peak regions. Genes enriched in the regenerating epicardium included *hypoxia-inducible factor 1ab* (*hif1ab*) (FDR=0.0052), shown to be required for zebrafish heart regeneration (Jopling, et al., 2012); the zinc finger and BTB domain-containing factor *zbtb7a* (FDR=3*10^-9^), a regulator of genes associated with heart failure following myocardial infarction (Niu, et al., 2019), and *tbx2b* (FDR=1*10^-6^), a known regulator of atrioventricular canal formation (Chi, et al., 2008). We also found several TFs which, similarly to *tcf21* and *tbx18*, were expressed in both the larval and the regenerating epicardium. One of them was *signal transducer and activator of transcription 3* (*stat3*), which in humans has cardio-protective functions (Zouein, et al., 2015) and promotes cardiomyocyte proliferation following zebrafish heart injury (Fang, et al., 2013).

**Figure 5.**
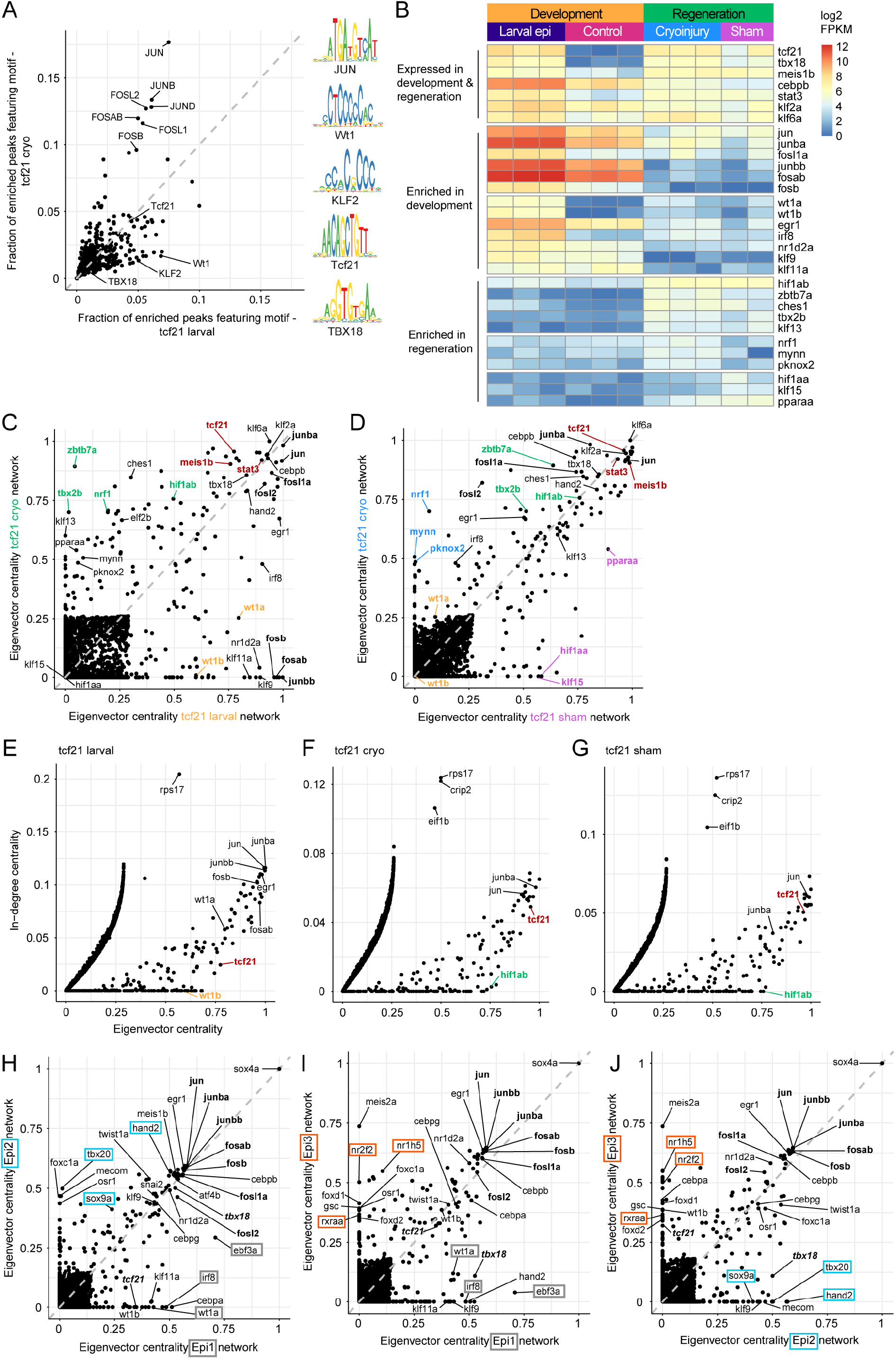
Gene regulatory networks identify central transcriptional regulators in the developing and regenerating zebrafish epicardium. (A) Quantification of motif presence in differentially accessible peaks in tcf21 larval versus tcf21 cryo. Shown is the fraction of peaks featuring a motif. Equal motif presence in both peak sets is indicated by a grey dotted line. Selected motif logos are shown on the right-hand side. (B) Bulk expression of transcription factors in the zebrafish heart at 5dpf (Development), as well as in the epicardium at 3dpi (Regeneration, Cryo) and at 3dps (Regeneration, Sham). Shown are log2 transformed FPKM values. (C,D) Comparison of eigenvector centrality in tcf21 larval and tcf21 cryo networks (C), and in tcf21 sham and tcf21 cryo networks (D). TFs with enriched centrality in cryoinjured epicardium are shown in blue, TFs with enriched centrality in sham-treated epicardium in purple. (E-G) In-degree centrality (y-axis) versus eigenvector centrality (x-axis) in tcf21 larval (E), tcf21 cryo (F) and tcf21 sham (G) sub-networks. (H-J) Comparison of eigenvector centrality in Epi1 and Epi2 networks (H), in Epi1 and Epi3 networks (I) and in Epi2 and Epi3 networks (J). tcf21 and tbx18 are labelled in bold italic letters. In H and I, factors with specific centrality in the Epi1 network are highlighted by a grey box. In H and J, factors with specific centrality in the Epi2 network are highlighted by a cyan box. In I and J, factors with specific centrality in the Epi3 network are highlighted by an orange box. In C,D,H-J, a grey dotted line indicates equal centrality. In C-G, marker regulators of both larval and adult epicardium are shown in red, marker regulators of larval epicardium are shown in yellow, marker regulators of adult epicardium are shown in green. In C-J, ap-1 components are labelled in bold letters. epi=epicardium. See also Figs. S3-5.

Building on these data, we constructed epicardial gene regulatory networks in both contexts to gain further insight into the transcriptional programmes driving the developing and regenerating epicardium. To this end, we combined chromatin accessibility and gene expression data using a recently developed computational framework (Xu, et al., 2021) (Figure S3B). We generated subsets of networks to retain only the strongest 1% of connections, yielding sub-networks of approximately 100,000 connections (Figure S3C). All sub-networks featured 200-250 TFs that acted as connection source nodes (n(larval epicardium)=230, n(cryoinjured epicardium)=216, n(sham-treated epicardium)=214), and a high number of target genes (n(larval epicardium)=1798, n(cryoinjured epicardium)=2445, n(sham-treated epicardium)=2422) (Figure S3D). To predict the connectivity within the network and thereby the regulatory importance of TFs, we computed the eigenvector centrality of each factor across networks (Nepusz, 2006). Comparing between the larval and the cryoinjured epicardial networks, we found that TFs such as tcf21, stat3 and meis1b, as well as the AP-1 components jun, junba, fosl1a and fosl2 possessed high centrality values in both networks, indicating they might play a role in epicardial transcriptional regulation during both development and regeneration (Figure 5C). In contrast, TFs such as wt1a, wt1b, junbb, fosab and fosb were much more central in the developing epicardial network than in the regenerating network, while TFs including hif1ab, zbtb7a, tbx2b and nrf1 showed the opposite trend and were used during regeneration. To further identify TFs that might play a regulatory role specifically during heart regeneration, we compared centrality between cryoinjured and sham-treated epicardial networks (Figure 5D). Many factors central in the cryoinjured epicardial network, including tcf21 and hif1ab, were similarly central in the sham-treated counterpart. However, TFs such as nrf1, mynn and pknox2 featured high centrality specifically in the cryoinjured epicardial network, while klf15, hif1aa, pparaa and others showed higher centrality in the larval network. We then asked if TFs with high network centrality were also positioned at the top of the regulatory TF hierarchy. To this end, we compared eigenvector centrality to in-degree centrality (IDC, a measure of the number of incoming network connections) (Nepusz, 2006), allowing us to characterise TFs based on their network connectivity, as well as the amount of regulatory input they received. We found that wt1b (IDC=0), but not AP-1 components such as jun (IDC=0.116) and fosab (IDC=0.083), were among the TFs predicted to reside at the top of the regulatory hierarchy in the larval epicardium (Figure 5E). In contrast, hif1ab was predicted to reside at the top of the hierarchies in the adult epicardial conditions (IDC(cryoinjured)=0.003, IDC(sham-treated)=0) (Figures 5F,G). Interestingly, tcf21 was predicted to receive a lower amount of regulatory input in the larval epicardium than in the adult activated epicardium (IDC(larval)=0.024, IDC(cryoinjured)=0.049, IDC(sham-treated)=0.051), suggesting tcf21 might move down the epicardial regulatory hierarchy after heart development and during heart regeneration may play a more downstream role.

We next focused on the larval epicardial regulatory networks to investigate the importance of TFs expressed in the developing Epi1-3 subpopulations (Figure S4A). We analysed the most robust 1% of connections (Figures S4B,C) and identified TFs whose centrality was specific to each subpopulation. We found that TFs such as wt1a, the interferon regulatory factor irf8 and ebf3a had the increased centrality specifically in Epi1 (Figures 5H,I). In contrast, Epi2 featured high centrality of TFs known to affect cardiac outflow tract formation such as tbx20 (Takeuchi, et al., 2005) and hand2 (Schindler, et al., 2014), as well as the mesenchymal cell marker sox9a (Figures 5H,J). On the other hand, TFs with specific centrality in Epi3 included the retinoic acid receptor rxraa, and the nuclear receptors nr1h5 and nr2f2 (Figures 5I,J). Our analysis also predicted that wt1b and tcf21 are positioned at the top of the regulatory hierarchies in Epi1 and Epi3 (Figures S4D,E), matching the results in the larval epicardial network (Figure 5E). Our analysis further predicted ebf3a to be a top-level regulator in Epi1 (Figure S4D), sox9a to be at the top level of the regulatory hierarchy in Epi2 (Figure S4F), and rxraa in Epi3 (Figure S4E).

In summary, our gene regulatory network analysis predicts specific TFs that play a central role in regulating epicardial gene expression (See Figures S5A,B for a complete overview). We highlight TFs inferred to be central during both heart development and regeneration, such as tcf21, meis1b and stat3, and TFs predicted to be of specific importance in either setting, such as wt1a and wt1b in the developing epicardium and hif1ab and nrf1 in the regenerating epicardium.

### Specific sets of epicardial upstream regulators drive post-injury transcriptional re-activation of developmental genes

After defining TFs that are *bona fide* epicardial regulators during development and regeneration, we asked whether there might be differential epicardial regulation even at the level of the small set of developmental genes that are re-activated upon injury (as suggested in Figure 4E). To this end, we compared the scaled strength of network connections between the commonly-expressed genes (Table S1) and the TFs tcf21, meis1b, stat3 (predicted to be central in the epicardium during heart development and regeneration), wt1a, wt1b (more central in the developing epicardium), as well as hif1ab and nrf1 (more central in the regenerating epicardium) (Figures 6A,B). We found that tcf21, meis1b and stat3 featured high connection scores in both the larval and the cryoinjured epicardial network, in line with the high eigenvector centralities these regulators featured in both networks (Figure 5E). However, wt1a and wt1b mainly featured stronger connections in the larval epicardial network, corroborating the finding that these TFs showed much higher centralities in the larval than in the cryoinjured epicardial network. Conversely, hif1ab and nrf1 were more central in the cryoinjured epicardial network, and their connections to epicardial genes commonly expressed during heart development and regeneration generally showed an increase in strength in the cryoinjured epicardial network. This analysis suggested that even for transcripts present during both development and regeneration, divergent sets of upstream TFs regulate their expression.

**Figure 6.**
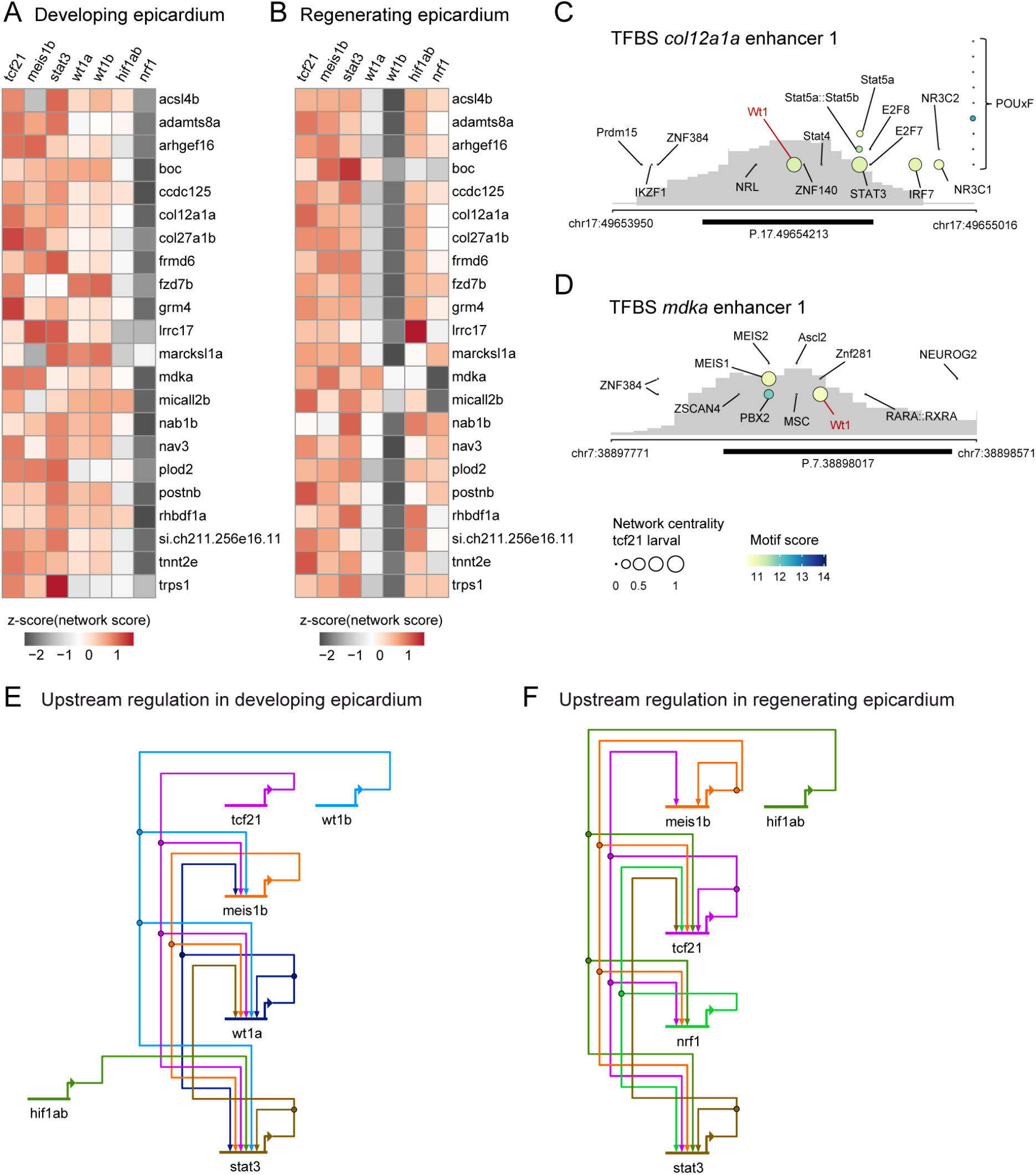
Differential transcriptomic regulation of genes expressed in both developing and regenerating zebrafish epicardium. (A,B) Row-wise z-scores of network connection scores between selected transcription factors and injury-reactivated developmental epicardial genes in the larval epicardium (A) and cryoinjured epicardium (B). (C) Transcription factor binding sites within the *col12a1a* enhancer 1 sequence. (D) Transcription factor binding sites within the *mdka* enhancer 1 sequence. In C and D, a Wt1 binding site is highlighted in red. Bubble size indicates the eigenvector centrality of the factor in the tcf21 larval sub-network (In case of duplicated zebrafish genes, the centrality of the higher-scoring paralogue is indicated). The quality of the binding sites is shown as colour-code. Accessibility of the endogenous genomic region in the tcf21 larval condition is underlaid in grey. A black bar indicates the position of the endogenous peak region. (E) Network schematic depicting predicted upstream transcriptional regulation between selected transcription factors in the tcf21 larval sub-network. (F) Network schematic depicting upstream regulation in the tcf21 cryo sub-network. In E and F, all interactions are activating. TFBS=transcription factor binding sites. See also Fig. S6.

We next assessed which TFs might drive *in vivo* larval cardiac activities of col12a1a enhancer 1 and mdka enhancer 1; regulatory regions with enriched accessibility in the larval epicardium. In both enhancer sequences, we identified a prominent Wt1 binding motif (col12a1a_e1: ATGTGTGGGAGGAA, reverse strand, motif score=11.1, mdka_e1: CTCCTCCCCCATGC, forward strand, motif score=10.8) (Figures 6C,D). In addition, in the col12a1a sequence, we identified a STAT3 motif (TTTCCAAGAAG, reverse strand, motif score=11.1), while the mdka enhancer 1 sequence also featured a MEIS1 motif (AGTGATTTATGAC, reverse strand, motif score=10.9). In both enhancer sequences, the chromatin surrounding the motif locations was accessible in the larval epicardium, corroborating the possibility of TF binding. This analysis inferred that TFs identified as *bona fide* central regulators of epicardial development directly activate larval-specific enhancer elements associated with genes commonly expressed in both the developing and regenerating epicardium.

Next, we focused on the identified *bona fide* epicardial upstream regulators and investigated their connections in the gene regulatory networks driving epicardial development and regeneration. For this, we visualised the strongest connections between tcf21, meis1b, stat3, wt1a, wt1b, hif1ab and nrf1 in both networks (Figures 6E,F) and found striking differences. While wt1b was positioned at the top of the regulatory hierarchy in the larval epicardial condition, connecting to wt1a, meis1b and stat3, it was absent from the cryoinjured epicardial condition. Its paralogue wt1a, involved in *bona fide* upstream regulation of the larval epicardium, was also absent from the *bona fide* regulatory circuits controlling the cryoinjured epicardium. In contrast, hif1ab only played a marginal role in the larval epicardial regulation but was well connected in the epicardial regulation activated upon injury. For example, hif1ab connected to nrf1, another regulator exclusively featuring in the upstream regulatory apparatus activated during regeneration. Matching the lower IDC in the larval epicardial network (Figure 5E), tcf21 resided at the top of the TF hierarchy regulating the developing epicardium. However, in the *bona fide* regulatory network of the cryoinjured epicardium, tcf21 received regulatory input from multiple sources, including itself. This might reflect a need to quickly upregulate *tcf21* expression following injury, contrasting with a more steady-state expression in the developing epicardium. The TF with the most comparable position in the regulatory networks was stat3, located at the bottom of the regulatory hierarchy in both larval and cryoinjured epicardial networks. Thus, we identified marked differences in the predicted upstream transcriptional regulation of the small set of genes commonly expressed during both epicardial development and regeneration.

We next focused our analysis on TF motifs in regulatory regions associated with gene markers enriched in specific larval epicardial subpopulations. Epi1 associated ppfibp1a enhancer 1 showed *in vivo* activity in epicardial cells (Figure 3I) and featured two TWIST1 motifs (CCTCCAGATGTGA, motif score=10.1 and TGGCCAGATGTGG, motif score=9.7) (Figure S6A), whereas Epi1 associated sema3e enhancer 12, which also showed *in vivo* activity in epicardial cells (Figure S2D), featured multiple KLF9 motifs (tgtgtgtgtgtgtgtg, motif score=9.6) (Figure S6B). Therefore, these Epi1-associated enhancers might be bound by twist1a and klf9, respectively, TFs with high centrality in Epi1 (Figures 5G,H). The Epi2 marker gene *loxa* and its putative enhancer 1, which showed *in vivo* activity in BA epicardial cells (Figure 3G) featured FOXC1 (TAAATAAATAA, motif score=9.6), CEBPG (TTCATTGCATCATGT, motif score=11.4) and ATF4 (TCATTGCATCATGT, motif score=12) motifs (Figure S6C), whereas Epi2 marker gene *elnb* and its putative enhancer 2, which also showed *in vivo* activity in BA epicardial cells (Figure S2B), featured SNAI2 (AGACACCTGTCTT, motif score=10.1) and MEIS1 (TACATAAATCACG, motif score=10.1) motifs (Figure S6D). Accordingly, the TFs foxc1a, cebpg, atf4b, snai2 and meis1b all showed high centrality in our newly-generated Epi2 network (Figures 5G,I). Thus, these TFs are putative top upstream drivers of the cell-specific activity reported by the newly identified developmental epicardial enhancers, *loxa*-enhancer 1 and *elnb-*enhancer 2.

Finally, we compared the inferred upstream regulation embedded in Epi1 and Epi2 subpopulation networks. We found that tcf21 and wt1b were positioned at the top of the TF hierarchy of Epi1, whereas TFs such as tbx18 and irf8 resided further down the hierarchy (Figure S6E). In the *bona fide* upstream regulatory network of Epi2, cebpg and snai2 were positioned higher up the hierarchy than TFs such as meis1b and tbx18 (Figure S6F). Therefore, our *in silico* analysis yielded new insights into how subpopulation-specific gene expression might be regulated in the developing epicardium.

## Discussion

The epicardium is an essential source of cardiovascular derivatives and mitogenic signals during heart development and regeneration, which has emerged as a potential target in the treatment of cardiovascular disease. Therefore, understanding the underlying genetic programmes orchestrating epicardial development and its response to injury is critical for successful therapeutic manipulation of the epicardium.

It is generally accepted that the transcriptional programmes employed during embryonic development are often repurposed during regeneration, yet the genetic mechanisms underlying such programmes remain elusive. In this study, we performed an unbiased genome-wide comparison of the epicardial gene programmes acting during zebrafish epicardial development and following adult heart injury. Surprisingly, we found that the transcriptomic programmes deployed during regeneration are not a mere recapitulation of the transcriptional activity driving embryonic development. By combining chromatin accessibility and gene expression data analysis and integration of TF expression levels and TF motif occupancy in peaks differentially accessible in the developing and the regenerating epicardium, we built gene regulatory circuits that specifically govern each of these processes. Although TFs such as tcf21, stat3, meis1b, and the AP-1 components jun, junba, fosl1a and fosl2 showed high centrality values in both networks, we identified wt1a, wt1b, and the AP-1 subunits junbb, fosab and fosb as central regulators in the developing epicardial gene regulatory network. Notably, expression of the AP-1 subunits junbb, fosab and fosb was preferentially found in the developing epicardial network, suggesting that specific AP-1 subunit compositions may be required to activate the embryonic epicardial transcriptional programme. Furthermore, our analysis predicted wt1b, but not the AP-1 components, to be at the top of the regulatory hierarchy during epicardial development. Conversely, the regenerating epicardial gene regulatory network featured TFs such as hif1ab, zbtb7a, tbx2b and nrf1 as putative central regulators, with hif1ab predicted to reside at the top of the regulatory hierarchy in the cryoinjured epicardium. We also found evidence that even for the group of transcripts shared by the embryonic and the regenerating epicardium, the underlying genetic programmes driving their expression are nevertheless distinct. Here, tcf21, meis1b and stat3 featured high connection scores in both the developing and the regeneration epicardial network, while wt1a and wt1b mainly featured stronger connections in the larval epicardial network. Conversely, hif1ab and nrf1 connections to epicardial genes commonly expressed during heart development and regeneration showed stronger connections in the regenerating epicardial network.

Further analysis revealed additional differences between the gene regulatory networks driving epicardial development and regeneration. Wt1b exclusively featured at the top of the regulatory hierarchy in the larval epicardial network, connecting to wt1a, meis1b and stat3. In contrast, our analysis showed hif1ab to be a central hub in the regeneration network, connecting, for example, to nrf1, a TF exclusively featured in the regenerating epicardial network. Notably, tcf21 positioning in each of the networks was strikingly different: while residing at the top of the developing TF hierarchy, tcf21 received regulatory input from multiple sources, including itself, in the regenerating epicardial network. We observed a similar finding when comparing the number of regulatory inputs that tcf21 received in the developing- and regeneration-specific transcriptional networks. This suggests that regulation of *tcf21* expression in the regenerating epicardium allows for multiple upstream inputs, which can potentially facilitate a rapid onset of expression upon injury, contrasting to the slower genomic regulatory blueprint deployed during development.

This study has also uncovered novel enhancer regions that specifically drive the transcriptional heterogeneity underlying epicardial development. We showed that putative regulatory regions associated with previously identified Epi1-3 epicardial subpopulations (Weinberger, et al., 2020) drove *in vivo* gene expression in a cell-type-specific manner. The newly identified Epi1-associated enhancers ppfibp1a_e1 and sema3e_e12 reported epicardial activity in cells which strongly resembled those of Epi1 origin. loxa_e1 and elnb_e2 enhancers drove specific reporter activity in the epicardial cells of the outflow tract (Weinberger, et al., 2020), further validating the Epi2-related specificity of the newly identified regulatory sequences. Building subpopulation-specific gene regulatory networks revealed multiple regulatory programmes active in the developing epicardium: Epi1 network featured wt1a, irf8 and klf11a as core upstream regulators, while Epi2 featured high centrality factors tbx20 and hand2, known to affect cardiac outflow tract formation (Xia, et al., 2019; Takeuchi, et al., 2005), as well as sox9a, as its top upstream inputs. Conversely, Epi3 network featured rxraa and the nuclear receptors nr1h5 and nr2f2 as high centrality TFs.

As noted above, tissue regeneration is often associated with the reactivation of embryonic gene programmes. That said, regeneration-specific gene programmes have been suggested in frog tadpole tail (Aztekin, et al., 2019), sea star limb (Oulhen, et al., 2016), axolotl limb (Gerber, et al., 2018), mouse digit-tip (Storer, et al., 2020) and crustacean leg (Sinigaglia, et al., 2021) regeneration studies. In addition, a few studies identified several cardiac injury TREEs (Begeman, et al., 2020; Wang, et al., 2020a; Goldman, et al., 2017; Pfefferli and Jazwinska, 2017; Kang, et al., 2016). To date, no study has systematically compared developmental and regenerative gene programmes and their regulation in a single cell type. Our work is the first to undertake a genome-wide approach based on epicardial gene expression and chromatin accessibility profiles to build, compare and contrast gene regulatory networks specifically driving epicardial genetic programmes during embryonic development and regeneration. Here we provide insight into the differential regulation of each of these processes and propose that the slower process of epicardial development involves a pre-planned series of coordinated events. In contrast, injury and regeneration entail a rapid response mode, and, as such, the positioning, utilisation and occupancy of regulatory elements are different in each of these processes. A deep understanding of the gene regulatory mechanisms underlying dynamic changes in epicardial gene expression is key to potentially develop therapeutics that can induce regeneration of the non-regenerative mammalian heart.

## Acknowledgments

We would like to thank the Biomedical Services Unit for fish husbandry as well as MRC WIMM Flow Cytometry Facility, MRC WIMM Sequencing Facility, MRC WIMM Single Cell Core Facility and the Wolfson Imaging Centre Oxford for excellent services. We thank Seda Ates and Jasmina Kuburic for technical assistance. This work was supported by BHF grants RG/13/9/303269 and CH/11/1/28798 to PRR, MW, and FCS, Oxford BHF Centre of Research Excellence Fellowship RE/13/1/30181 award to FCS; and Wellcome Trust Senior Research Fellowship (215615/Z/19/Z) to TSS.

## Author Contributions

Conceptualisation, MW, FCS, TSS and PRR; Methodology, MW and FCS; Investigation, MW and FCS; Computational analysis and Data Curation, MW; Writing – Original Draft, MW and FCS; Writing – Review & Editing, MW, FCS, TSS and PRR; Supervision, TSS and PRR; Funding Acquisition, TSS and PRR.

## Materials and Methods

### Materials availability

Further information and requests for resources and reagents should be directed to and will be fulfilled by Paul Riley. Requests for zebrafish transgenic lines should be directed to Tatjana Sauka-Spengler.

### Experimental Model and Subject Details

For this study, both females and males of transgenic and wildtype zebrafish strains were used. Animals used for breeding were between 3 and 24 months old. Zebrafish larvae that were used for experiments were raised to an age of 5 days post fertilisation (dpf). Larvae were euthanised and analysed shortly before reaching an age of 5dpf (free-feeding) during all experiments for which the experimental timepoint is stated as “5dpf” in text or figures. Fish were kept at a 14 hours light, 10 hours dark cycle and fed four times a day. All animal experiments were performed under a Home Office Licence according to the Animals Scientific Procedures Act 1986, UK, and approved by the local ethics committee.

### Zebrafish Lines

Published transgenic reporter lines used in this study were: *TgBAC(tcf21:H2B-Dendra2)^ox182^* (Weinberger, et al., 2020), *TgBAC(tbx18:myr-Citrine)^ox185^* (Weinberger, et al., 2020) and *Tg(βactin:Avi-Cerulean-RanGap)^ct700a^* (Trinh, et al., 2017).

### Generation of transgenic zebrafish lines

To generate *TgBAC(tcf21:BirA-2a-mCherry)^ox143^*, we used a BAC recombineering approach (Bussmann and Schulte-Merker, 2011). Briefly, a BirA-2A-Cherry-SV40pA-FRT-Kan-FRT cassette was PCR-amplified using Herculase II fusion DNA polymerase (Agilent Technologies) and recombined into the first coding exon of *tcf21* within the DKEYP 79F12 BAC clone as previously described (Weinberger, et al., 2020). In a second recombination step, an iTol2-Ampicillin cassette (provided by Prof. Kawakami, National Institute of Genetics, Mishima, Japan) was introduced into the BAC backbone as previously published (Bussmann and Schulte-Merker, 2011). Wild-type embryos were injected at the one-cell stage with 200 ng/μl of purified BAC DNA and 100 ng/μl *tol2* transposase mRNA. Putative founders were outcrossed to wild-type fish and offspring screened for Cherry expression, in combination with PCR amplification of the transgene when expression levels were low. *TgBAC(tcf21:BirA-2A-Cherry;βactin:Avi-Cerulean-RanGap)^ox144^* was generated by crossing *TgBAC(tcf21:BirA-2a-mCherry)^ox143^* (generated in this study) to *Tg(βactin:Avi-Cerulean-RanGap)^ct700a^* (Trinh, et al., 2017).

### Hybridisation Chain Reaction

Wildtype larvae were euthanised using tricaine methanesulfonate (MS-222) and fixed in 4% paraformaldehyde (PFA) overnight at 4°C. Subsequently, larvae were stored at -20°C in methanol. Hybridisation chain reaction (HCR) v3.0 (Choi, et al., 2018) was performed following a protocol by Choi et al. Briefly, larvae were permeabilized using 30μg/ml proteinase K for 45 minutes at room temperature, post-fixed in 4% PFA and incubated overnight at 37°C in 30% probe hybridisation buffer containing 2pmol of each probe mixture. Excess probes were washed off with 30% probe wash buffer at 37°C and 5X Sodium Chloride Sodium Citrate/0.1% Tween 20 (SSCT) at room temperature and larvae were incubated overnight at room temperature in amplification buffer containing 15pmol of each fluorescently labelled hairpin. Following HCR, larvae were incubated with Hoechst reagent (1:1000, 5XSSCT) for 30 minutes at room temperature.

Adult injured and sham-control hearts were collected and fixed in 4% paraformaldehyde (PFA) for 2h at room temperature. OCT-embedded hearts were cryosectioned and 10μm sections were then washed in DEPC-treated water and permeabilised for 10 minutes using PBS/0.1% Tween 20. Staining was performed as previously described (Choi, et al., 2018), with the following adaptations: sections were pre-hybridised with 30% probe hybridization buffer for 10 min at 37 °C in a humidified chamber. Sections were then incubated with 1.6 pmol of each DNA probe (Molecular Instruments) diluted in hybridization buffer, covered with a cover slip and incubated overnight at 37 °C in a humidified chamber. Sections were washed in 30% probe wash buffer and 5X Sodium Chloride Sodium Citrate/0.1% Tween 20 (SSCT) at room temperature and then incubated for 30 minutes at room temperature in amplification buffer. Hairpins were incubated at final concentration of 6 pmol each (amplifier B1-Alexa488, amplifier B3-Alexa594, and amplifier B4-Alexa546; Molecular Instruments), overnight at room temperature in a dark humidified chamber. Excess hairpins were washed in 5X SSCT and nuclei stained with Hoechst reagent (1:1000, 5XSSCT) for 10 minutes at room temperature.

Probe sequences were designed by the manufacturer probe (Molecular Instruments), probe sets used were: dr_tcf21 (amplifier B1), col12a1a_dr (amplifier B3), mdka_dr (amplifier B4), plod2_dr (amplifier B3), postnb_dr (amplifier B4). Images were obtained using a LSM780 confocal microscope (ZEISS) using 10x and 40x objectives. Contrast and brightness were adjusted separately for each colour channel.

### Heart Isolation, Dissociation and FAC-Sorting

Larvae were euthanised using MS-222 and larval hearts were isolated following a published protocol (Burns and MacRae, 2006), using a 21-gauge needle for disruption. This procedure recovered around 50% of the larval hearts. Hearts were dissociated using 15mg/ml collagenase (C8176, Sigma Aldrich) in 0.05% trypsin solution at 30°C for 14mins. 7-AAD cell viability dye was used to exclude non-viable cells during FACS. 1000-2000 fluorescent cells and 1000-2000 non-fluorescent cells were purified from *TgBAC(tcf21:H2B-Dendra2)^ox182^* or *TgBAC(tbx18:myr-Citrine)^ox185^* (200 hearts). Cells sorted in separate FACS sessions were processed separately during RNA-seq or ATAC-seq library preparation.

Adult mCherry+ epicardial cells were FACS-isolated from *TgBAC(tcf21:BirA-Cherry)^ox143^* operated adult hearts (1 heart per sample). Prior to FACS, heart tissue was dissociated using 20 mg/ml collagenase in 0.05% Trypsin/0.53 mM EDTA/1xHBSS buffer to obtain single cell suspensions. Reaction was stopped in 10 mM HEPES/0.25% BSA/1xHBSS buffer and mCherry+ cells were sorted (FACSAria, BD Biosciences Fusion System). Cells sorted in separate FACS sessions (around 300 cells/sample) were processed separately during ATAC-seq library preparation.

### Adult zebrafish heart injury

Cardiac injuries were carried out in 4–12-month old zebrafish (Chablais, et al., 2011; Gonzalez-Rosa, et al., 2011). Briefly, cryoinjury was performed by application of a cryoprobe frozen with liquid nitrogen to the surface of the exposed ventricle until the probe was fully thawed, damaging approximately 20% of the ventricle. Exposing the ventricle, without injury, was performed for sham controls. Cryoinjured and sham-control hearts were harvested 3 days after injury/sham (3dpi and 3dps, respectively).

### Biotagged nuclei isolation

Biotagged nuclei were isolated as previously described (Simoes, et al., 2020; Trinh, et al., 2017). Briefly, *TgBAC(tcf21:BirA-Cherry;βactin:Avi-Cerulean-RanGap)^ox144^* operated adult hearts (n = 2 per sample) were washed and incubated on ice in hypotonic buffer H (20 mM HEPES, pH 7.9; 15 mM MgCl2; 10 mM KCl; 1 mM DTT; 1 X Complete protease inhibitor) for 30 min. Heart samples were transferred to a Dounce homogenizer (2 ml Kontes Glass Co, Vineland, NJ) and dissociated by 10 strokes with the loose fitting pestle A and incubated on ice for 5 min. Further dissociation was carried out by 10 strokes with tight fitting pestle B, performed every 5 min for 15 min. Nuclei were collected by centrifugation (2000 × g, 4 °C) and re-suspended in 1 ml of nuclei pulldown buffer NPB buffer (10 mM HEPES, pH 7.9; 40 mM NaCl; 90 mM KCl; 0.5 mM EDTA; 0.5 mM spermidine; 0.15 mM spermine; 1 mM dithiothreitol and 1 X Complete protease inhibitor). For nuclei purification, nuclei were incubated with 250 µg of M-280 streptavidin-coated dynabeads (Invitrogen) with rotation for 30 min at 4 °C. A flow-based system was used to capture the nuclei bound on the streptavidin beads. A 10 ml seriological pipette (VWR) attached to a 1 ml micropipette tip (Rainin reach pipet tip), both pre-treated with NPB+1% BSA for 30 min, was added to a MiniMACS separator magnet (OctoMACS Separator, Miltenyl Biotec). A two-way stopcock (Biorad) was connected to the end of the 1 ml micropipette tip via a piece of Tygon tubing (Fisher Scientific) and the flow-rate set to ∼0.75 ml/min. The nuclei beads suspension was diluted by addition of 9 ml of NPBt (NPB with 0.01% Triton X-100) and added to the slow-flow setup. The tip was subsequently removed from the stand and the nuclei-beads released from the tip by slowly pipetting 1 ml of NPBt in and out of the tip. The solution was then diluted again to 10 ml with NPBt and added again to the slow-flow setup. Nuclei-beads were eluted in 1 ml of NPBt as described above and the NPBt removed using a magnetic stand (DynaMag TM-2 magnet, Invitrogen). Nuclei-beads were then processed for RNA extraction.

### RNA extraction and library preparation for sequencing

1000 FACS-purified cells from *TgBAC(tcf21:H2B-Dendra2)^ox182^* larvae were processed using the SMART-SeqTm v4 UltraTm Low Input RNA Kit for Sequencing (Takara Clontech). Samples were lysed and poly-adenylated RNA reverse transcribed via SMARTScribe Reverse Transcriptase. Following reverse transcription, cDNA was amplified with SeqAmp DNA Polymerase, using 11 PCR cycles. Amplified cDNA was purified using Agencourt AMPure XP beads (Beckman Coulter). The purified cDNA was quantified using Qubit Fluorometric Quantitation (ThermoFisher) and the quality was verified on an Agilent 2100 Bioanalyzer (Agilent Technologies). Final sequencing libraries were prepared using the Nextera XT DNA Library Preparation Kit (Illumina). 1ng cDNA was used as input, samples were tagmented for 5 minutes and 10 seconds and amplified using 12 PCR cycles. The libraries were quantified using Qubit Fluorometric Quantitation (ThermoFisher) and their quality checked on a 2200 TapeStation system (Agilent). All cDNA libraries were pooled and sequenced to a depth of 355 million reads on a NextSeq500 machine (Illumina, 150 Cycle High Output Kit).

Total nuclear RNA extraction and DNAse treatment of *TgBAC(tcf21:BirA-Cherry;βactin:Avi-Cerulean-RanGap)^ox144^* adult-derived epicardial nuclei were carried out using the RNAqueous Micro Kit (Life Technologies) according to manufacturer’s instructions. RNA integrity was checked with an RNA pico chip (Agilent Technologies) using the Agilent 2100 Bioanalyzer (Agilent Technologies). cDNA was synthesized and amplified from 100 pg–300 pg of input RNA using SMART-seqTMv4 Ultra Low input RNA kit (Takara Clontech). Sequencing libraries were prepared using the Nextera XT DNA library preparation kit (Illumina). Next Generation Sequencing was performed on a NextSeq500 platform using a NextSeqTM500 150-cycle High Output Kit) (Illumina) to generate 80-basepair paired end reads.

### ATAC Sequencing

ATAC-seq was performed following a protocol modified from Buenrostro and colleagues (Buenrostro, et al., 2013). FACS purified cells from *TgBAC(tcf21:H2B-Dendra2)^ox182^* or *TgBAC(tbx18:myr-Citrine)^ox185^* larvae were pelleted at 600g for 7.5 minutes at 4°C, washed, pelleted, lysed and pelleted at 600g for 10 minutes at 4°C. Samples were then tagmented using 0.25µl Tn5 in 5µl TD buffer and 4.75µl H2O for 30 minutes at 37°C. Then, EDTA was added to a concentration of 50mM and samples were incubated for 30 minutes at 50°C. MgCl2 was added to a concentration of 50mM and 16µl of each sample were PCR amplified using 2x NEB Next HiFi PCR mix (NEB) and 16 PCR cycles.

FACS purified cells from *TgBAC(tcf21:BirA-2a-mCherry)^ox143^* adult hearts were pelleted at 500g for 5 minutes at 4°C, washed with ice cold 1X PBS buffer and pelleted again 500g for 5 minutes at 4°C. Cells were then lysed by gentle resuspension in cold lysis buffer (10 mMTris-HCl, pH 7.4; 10 mM NaCl; 3 mM MgCl2; 0.1% IGEPAL CA-630) and immediately spun down at 500xg for 10 minutes at 4°C. Samples were then tagmented using 0.25µl Tn5 in 5µl TD buffer and 4.75µl H2O for 20 minutes at 37°C. Samples were quenched by adding EDTA to a concentration of 50mM and incubated for 30 minutes at 50°C. MgCl2 was added to a final concentration of 50mM and 12.2µl of each tagmented sample was PCR amplified for 15 cycles using 2x NEB Next HiFi PCR mix (NEB). Amplified libraries were purified using the Qiagen PCR purification MinElute kit.

The quality of the ATAC libraries was checked on a 2200 TapeStation system (Agilent). Libraries were sequenced to a depth of approximately 50 million reads per sample (Illumina, 75 Cycle High Output Kits).

### Generation of enhancer reporter constructs

To generate loxa_e1, elnb_e2, ppfibp1a_e1, sema3e_e12, col12a1a_e1 and mdka_e1 reporter vectors, genomic sequences were amplified from wildtype DNA using Herculase II fusion DNA polymerase (Agilent Technologies). Sequence coordinates in GRCz10 and primers used for amplification are listed in Table S2. eGFP was replaced with mCherry in the pVC-Ds-E1b:eGFP-Ds plasmid (#102417, Addgene) (Chong-Morrison, et al., 2018). The plasmid was then linearised with NheI (NEB) (loxa_e1, elnb_e2) or with NheI and XhoI (NEB) (ppfibp1a_e1, sema3e_e12, col12a1a_e1 and mdka_e1) and amplified sequences were inserted using InFusion (InFusion HD Cloning kit, Clontech). Plasmid constructs were injected into one-cell zygotes and integrated into the genome via Ac-mediated recombination. Images of euthanised larvae were obtained using a LSM780 confocal microscope (ZEISS) and a 20x objective.

### Bioinformatic Processing

#### Published Datasets

Previously published scRNA-seq data (Weinberger, et al., 2020) was used to analyse TF expression in the larval epicardium and to construct larval epicardial networks (GEO accession number GSE121750). Raw and processed data generated in this study were submitted to GEO (accession number GSE178751).

#### RNA Sequencing Data Analysis

Transcriptomic data was mapped to the zebrafish reference genome (GRCz10-91) using the STAR gapped aligner (Dobin, et al., 2013). Duplicate reads were removed with samtools (v1.10) and reads were summarized using featureCounts (Liao, et al., 2014). Differential gene expression analysis and principal component analysis were performed in R (v4.0.3) using DESeq2 (v1.30.0) (Love, et al., 2014). Counts were transformed into FPKM expression values with the rpkm() command in the edgeR package (v3.32.1) (Robinson, et al., 2010) and heatmaps drawn using pheatmap (v1.0.12). Gene ontology term analysis was performed with the topGO package (v2.42.0) (Alexa and Rahnenfuhrer, 2021), using differentially expressed genes with an adjusted p-value below 0.001.

#### ATAC Sequencing Data Analysis

Sequencing reads were mapped to the zebrafish reference genome (GRCz10-91) using the bowtie aligner (v1.2.3) (Langmead, et al., 2009). Duplicate reads were removed with samtools (v1.10). BAM files were converted to bigWig format and accessibility tracks were visualised in the UCSC Genome Browser. BAM files were converted to BED format and peaks were called using MACS (v2.2.7.1, callpeak -B -f BED -g 1.37e09 --call-summits) (Zhang, et al., 2008). The Homer (v20201202) (Heinz, et al., 2010) script “annotatePeaks.pl” was used to annotate peaks to the closest expressed gene in each condition and to genomic features, to do TSS enrichment analysis and to compute CpG and GC content. For this, fasta and gtf files for GRCz10 were obtained from the UCSC Genome Browser (Kent, et al., 2002) website. The R package DiffBind (v3.0.13) (Stark and Brown, 2011) was then used to construct a consensus peak set across all samples, removing peaks contained in less than two samples and peaks overlapping repetitive elements in the genome, and resizing all peak widths to 500bp. Peak names were assigned based on genomic position as “P.[chromosome].[start position]”. Peak accessibility FPKM values were computed manually. DESeq2 (v1.30.0) was used for differential peak accessibility analysis. Gene ontology term analysis was performed with the R package topGO (v2.42.0), using genes that differentially accessible peaks with an adjusted p-value below 0.05 and a baseMean above 30 were annotated to.

#### Transcription Factor Binding Site Analysis

For analysis of TF binding in identified peak regions, TF motifs were identified across the consensus peak set, as well as in the putative enhancer sequences used for the *in vivo* enhancer activity assay, using the “gimmescan” command in the gimmemotifs package (van Heeringen and Veenstra, 2011) and the JASPAR2020 (v0.99.10) core vertebrates transcription factor motif database (Fornes, et al., 2020). A false positive rate of 0.05 (-f 0.05) was used as a cutoff when calling motifs across the consensus peak set. When calling motifs in the enhancer sequences, a cutoff score of 0.6 (-c 0.6) was used and motifs with a score below 10.5 (col12a1a_e1, mdka_e1) or 9.5 (loxa_e1, elnb_e2, ppfibp1a_e1, sema3e_e12) were subsequently excluded. The R chromVAR (v1.12.0) package (Schep, et al., 2017) was used cluster samples according to TF motif accessibility and to plot motif logos.

#### Gene Regulatory Networks

Gene regulatory networks were constructed using the Python package Ananse (v0.1.7). To construct the tcf21 larval network, input was extracted from tcf21 larval ATAC-seq and RNA-seq samples, tcf21 cryo samples were used for the tcf21 cryo network and tcf21 sham samples for the tcf21 sham network. For the Epi1 network, tcf21 larval and tbx18 larval ATAC-seq samples as well as pseudo-bulk data constructed from Epi1 cell scRNA-seq data was used as network input. For the Epi2 network, tbx18 larval ATAC-seq samples as well as pseudo-bulk data constructed from Epi2 cell scRNA-seq data was used. For the Epi3 network, tcf21 larval ATAC-seq samples as well as pseudo-bulk data constructed from Epi3 cell scRNA-seq data was used. As ATAC-seq input, a consensus peak set across samples of the relevant condition was constructed using DiffBind, and the mean FPKM accessibility values were computed. As RNA-seq input, mean TPM gene expression values were computed manually. Additionally, gene positions were extracted from the GRCz10 GTF file and stored in BED format. The JASPAR2020 (v0.99.10) core vertebrates transcription factor motif database was used to acquire TF binding motifs. “prob” values in the Ananse network output (termed “network scores” in Figure 6) were taken forward to indicate the strength of connections in the network. Ananse networks were subset to only contain connections with a strength above 0.99. The sub-networks were transformed into weighted directed graphs (“prob” values were used as weights) and the eigenvector centrality (evcent) computed using R igraph (v1.2.6). The in-degree centrality was computed via: degree(g, mode = “in”) / (vcount(g) -1), following a previous approach (Kamimoto, et al., 2020). Network graph schematics were plotted using BioTapestry (v7.1.2.0) (Longabaugh, et al., 2005), using network connections between the indicated TFs with a strength above 0.995.

#### Quantification and Statistical Analysis

Statistical details of experiments can be found in the figure legends, including p-values and numbers of samples analysed. Unless noted otherwise, adjusted p-values of below 0.05 were treated as significant. Plots were generated with ggplot2 (Wickham, 2016) in R. Some plots include box and whiskers plots (in the style of Tukey), indicating median and first/third quartiles.

