## Supplemental Figures, Supplemental Figure legends and Supplemental Tables for "Distinct epicardial gene regulatory programmes drive development and regeneration of the zebrafish heart"

### Supplemental Figures and Supplemental Figure legends

#### Figure S1

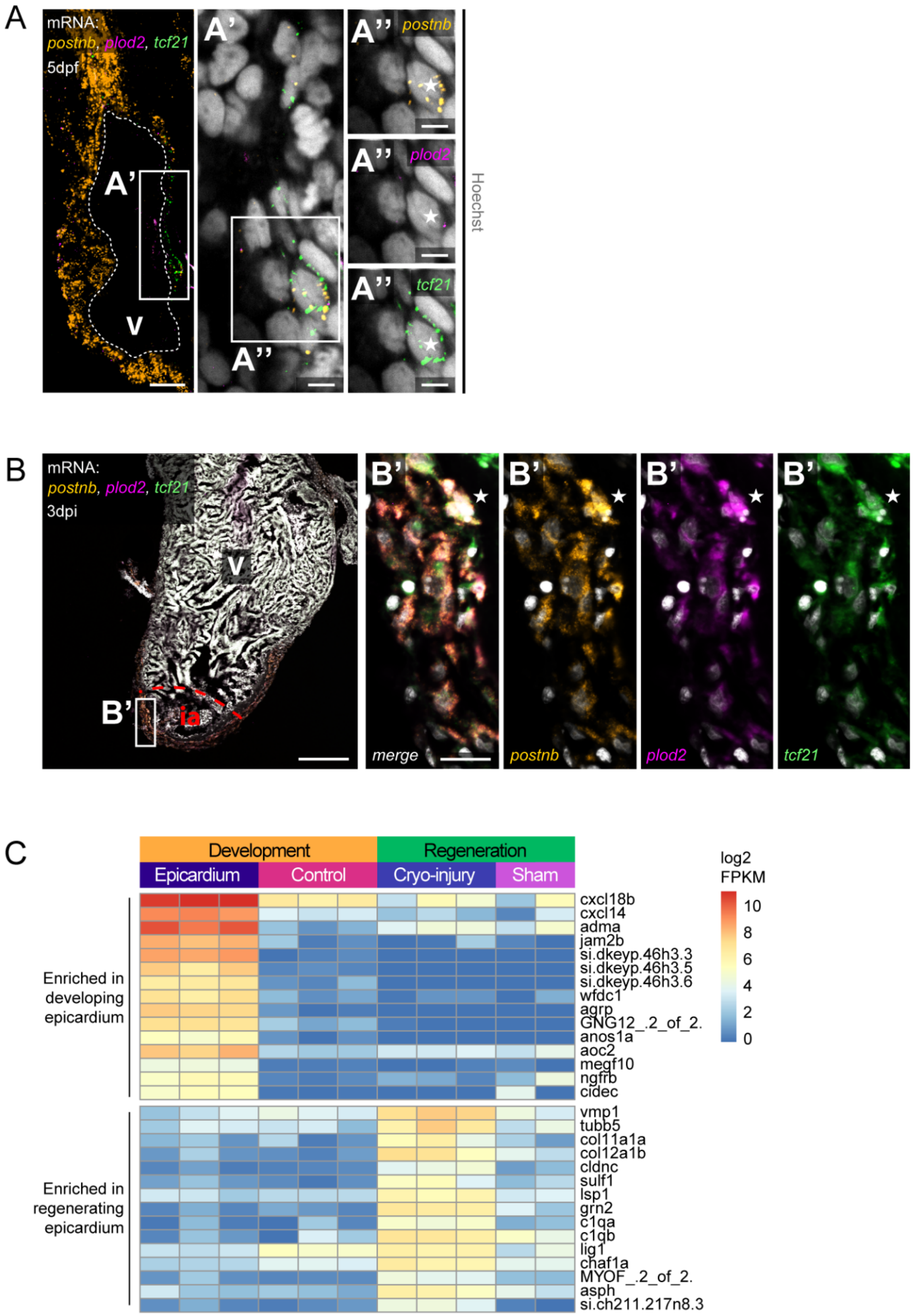

**Figure S1. Differential gene expression in developing and regenerating zebrafish epicardium.**

(A) mRNA staining of *postnb* (orange), *plod2* (magenta) and *tcf21* (green) in a 5dpf heart. (A‘,A‘‘) A nucleus (asterisk) in the epicardial region surrounded by *postnb*, *plod2* and *tcf21* transcripts. (B) mRNA staining of *postnb* (orange), *plod2* (magenta) and *tcf21* (green) in a 3dpi cryoinjured heart. ia=injury area. (B‘) A nucleus (asterisk) in the epicardial region surrounded by *postnb*, *plod2* and *tcf21* transcripts. Scale bars B: 100µm, A,B’: 20µm, A’,A’’: 5µm. Colour channels adjusted separately for brightness/contrast. A,B are single optical sections. (C) Heatmap showing differentially expressed genes in cryoinjured versus sham-injured adult epicardium. Shown are log2 transformed FPKM values. Related to Figure 1.

**Figure S2**

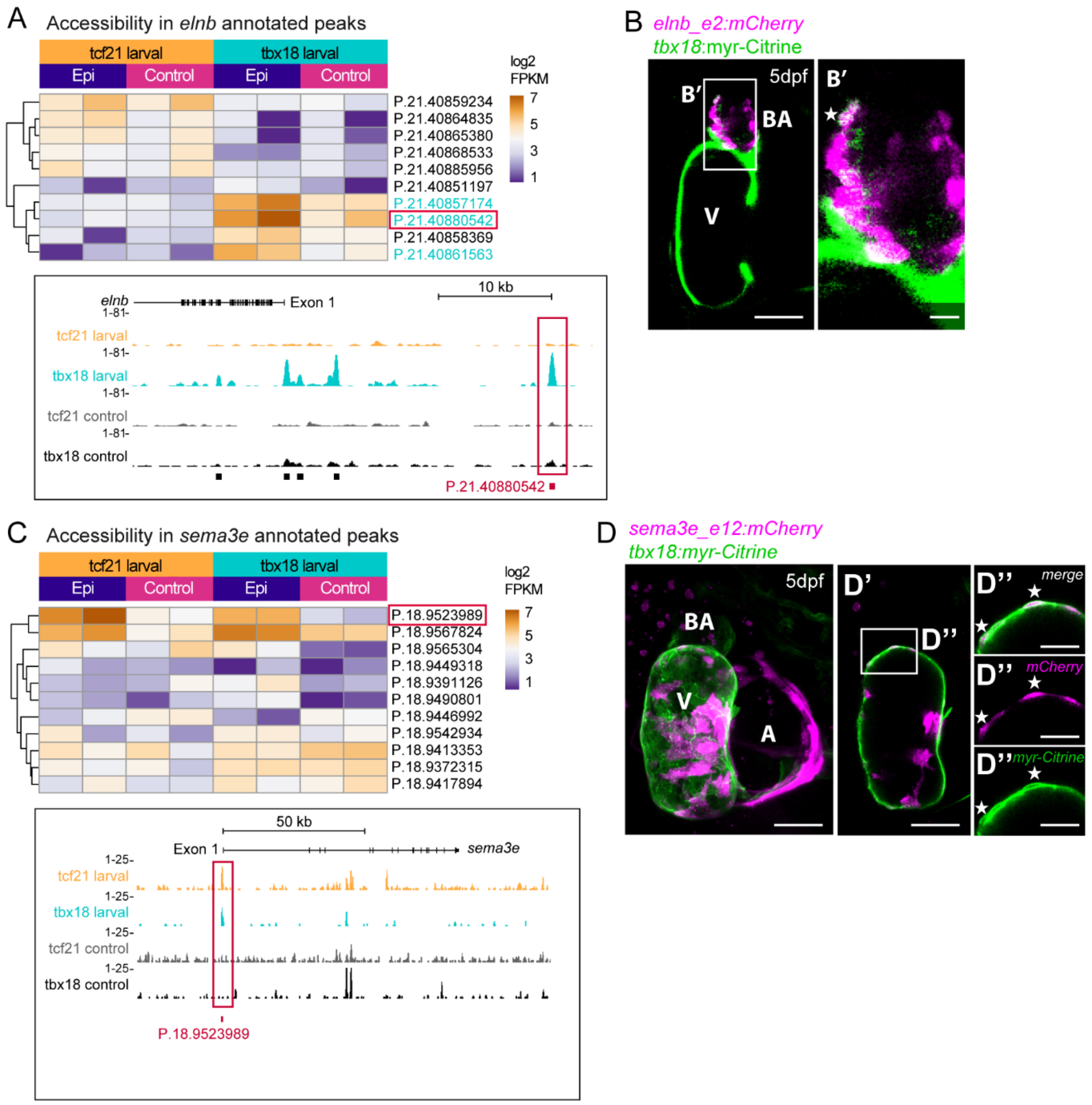


**Figure S2. *In vivo* enhancer activity of peak regions annotated to Epi2 and Epi1 marker genes.**

(A) Accessibility of peaks close to the Epi2 marker *elnb*. Cyan peak label colour indicates significant enrichment in tbx18 larval. A red frame indicates a peak analysed further in B. (B) elnb_e2 driven *in vivo* reporter expression (magenta) at 5dpf. Expression of *tbx18* is indicated by myr-Citrine fluorescence (green membranes). (B') Overlap of elnb_e2 activity and myr-Citrine in the BA (asterisk). (C) Accessibility of peaks close to *sema3e*. A red frame indicates a peak analysed further in D. (D) sema3e_e12 driven *in vivo* reporter expression (magenta) at 5dpf. Expression of *tbx18* is indicated by myr-Citrine fluorescence (green membranes). Close proximity of sema3e_e12 activity and myr-Citrine at the boundary of the ventricle (asterisks). (D‘) Single optical section from D. (D’’) mCherry,myr-Citrine$^{+}$ cells (asterisks) in the epicardial region. Scale bars in B,D,D’: 50µm, scale bar in B’: 10µm, scale bars in D’’: 20µm. V=ventricle, A=atrium, BA=bulbus arteriosus. Related to Figure 3.

### Figure S3

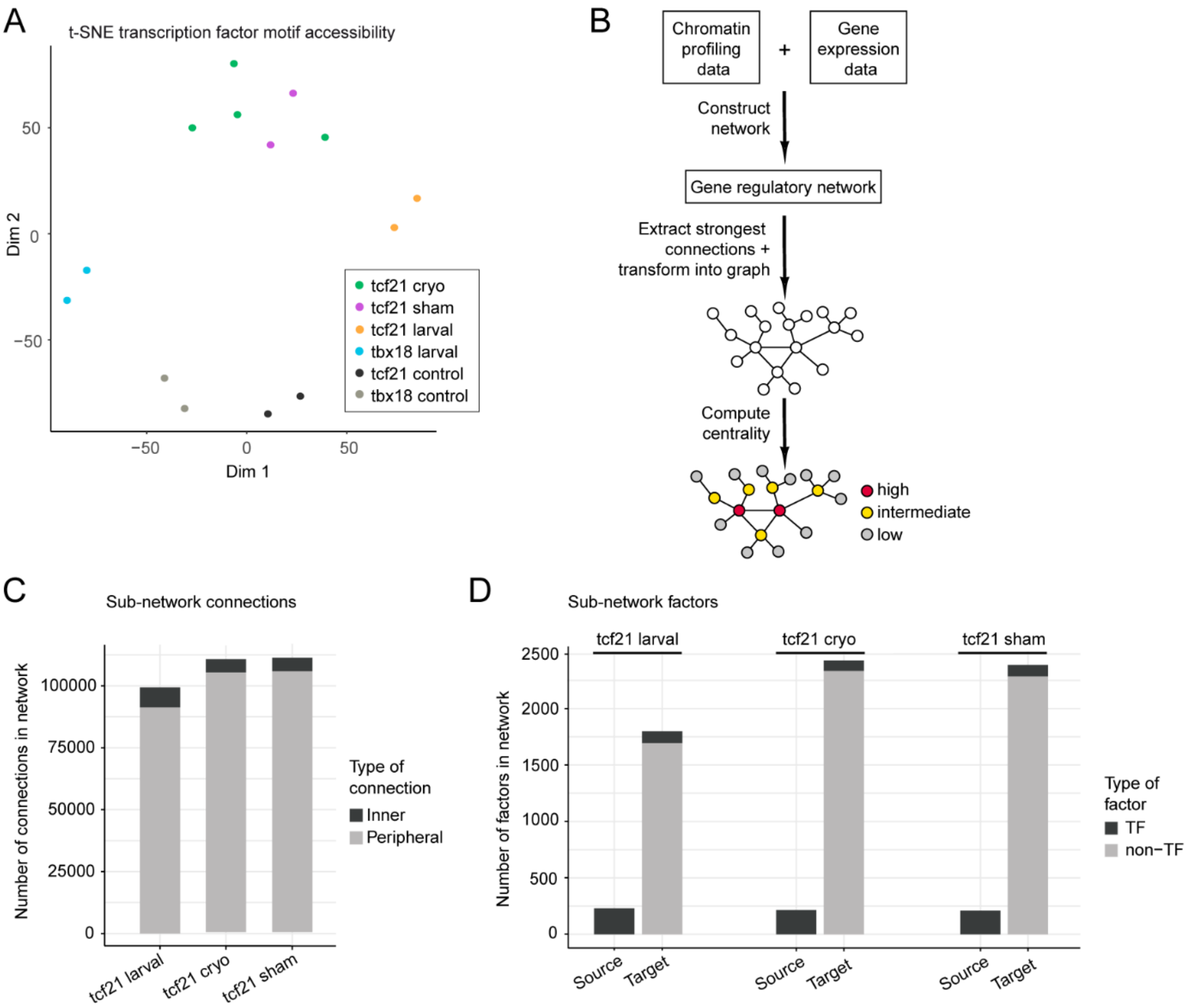


**Figure S3. Characteristics of gene regulatory networks in developing and regenerating epicardium**

(A) t-SNE clustering of samples according to transcription factor motif accessibility. (B) Schematic showing the workflow used to construct gene regulatory networks and to identify central regulators in those networks. (C) Number of connections in tcf21 larval, tcf21 cryo and tcf21 sham sub-networks. Shown are connections between TFs and factors featuring both incoming and outgoing connections (inner connections) and connections between TFs and factors featuring only incoming connections (peripheral connections). (D) Number of nodes (factors) in the epicardial sub-networks. Shown are the numbers of unique factors that act as source or as target of a connection. Target factors may be TFs or non-TFs. Related to Figure 5.

**Figure S4**

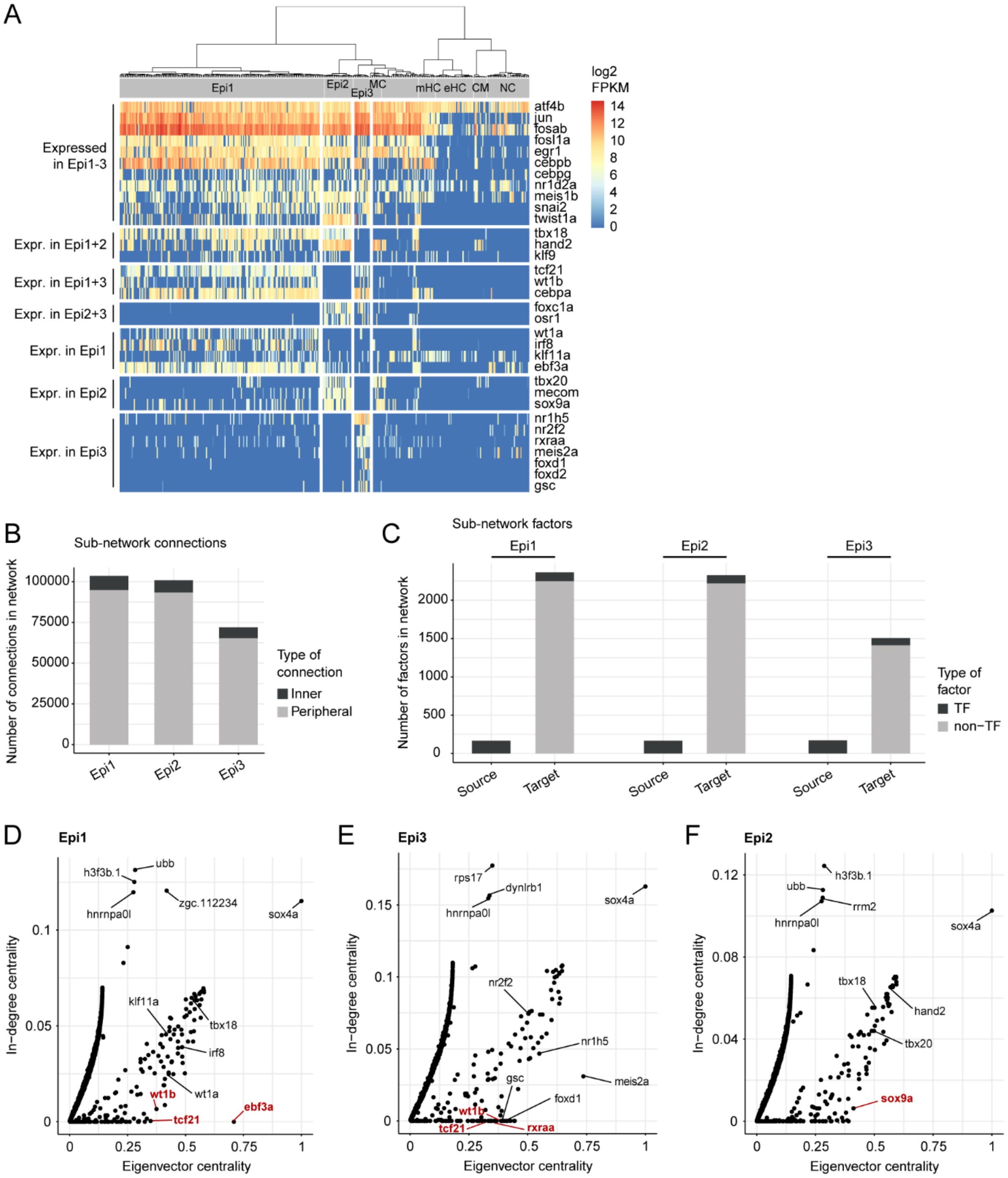


**Figure S4. Characteristics of gene regulatory networks of larval epicardial subpopulations**

(A) Heatmap showing scRNA-seq expression of transcription factors in the developing zebrafish heart at 5dpf. Shown are log2 transformed FPKM values. Epi=epicardium, MC=mesenchymal cells, mHC=myeloid haematopoietic cells, eHC=erythroid haematopoietic cells, CM=cardiomyocytes, NC=neural cells. (B) Number of connections in Epi1, Epi2 and Epi3 sub-networks. Shown are connections between TFs and factors featuring both incoming and outgoing connections (inner connections) and connections between TFs and factors featuring only incoming connections (peripheral connections). (C) Number of nodes (factors) in the larval epicardial sub-networks. Shown are the numbers of unique factors that act as source or as target of a connection. Target factors may be TFs or non-TFs. (D-F) In-degree centrality (y-axis) versus eigenvector centrality (x-axis) in Epi1 (D), Epi3 (E) and Epi2 (F) sub-networks. TFs mentioned in main text are highlighted in red. Related to Figure 5.

**Figure S5**

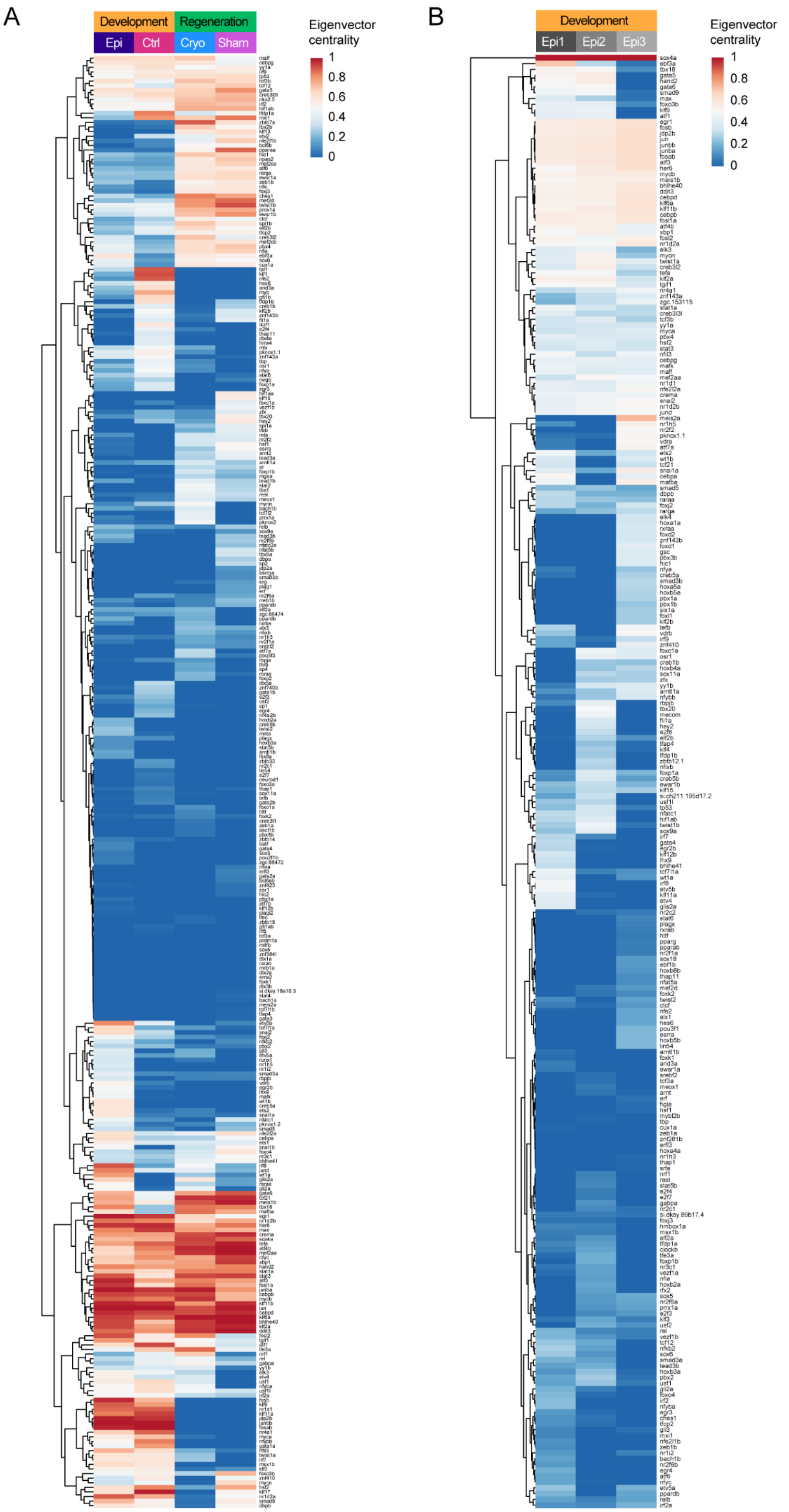
A
Development
Regeneration
Epi
Ctrl
Cryo
Sham
Eigenvector centrality
1
0.8
0.6
0.4
0.2
0
B
Development
Epi1
Epi2
Epi3
Eigenvector centrality
1
0.8
0.6
0.4
0.2
0

**Figure S5. Centrality in gene regulatory networks during zebrafish heart development and regeneration**

(A) Heatmap showing eigenvector centrality values of transcription factors in tcf21 larval (Development, Epi), tcf21 control (Development, Ctrl), tcf21 cryo (Regeneration, Cryo) and tcf21 sham (Regeneration, Sham) sub-networks. (B) Heatmap showing eigenvector centrality values of transcription factors in Epi1, Epi2 and Epi3 sub-networks. Related to Figure 5.

**Figure S6**

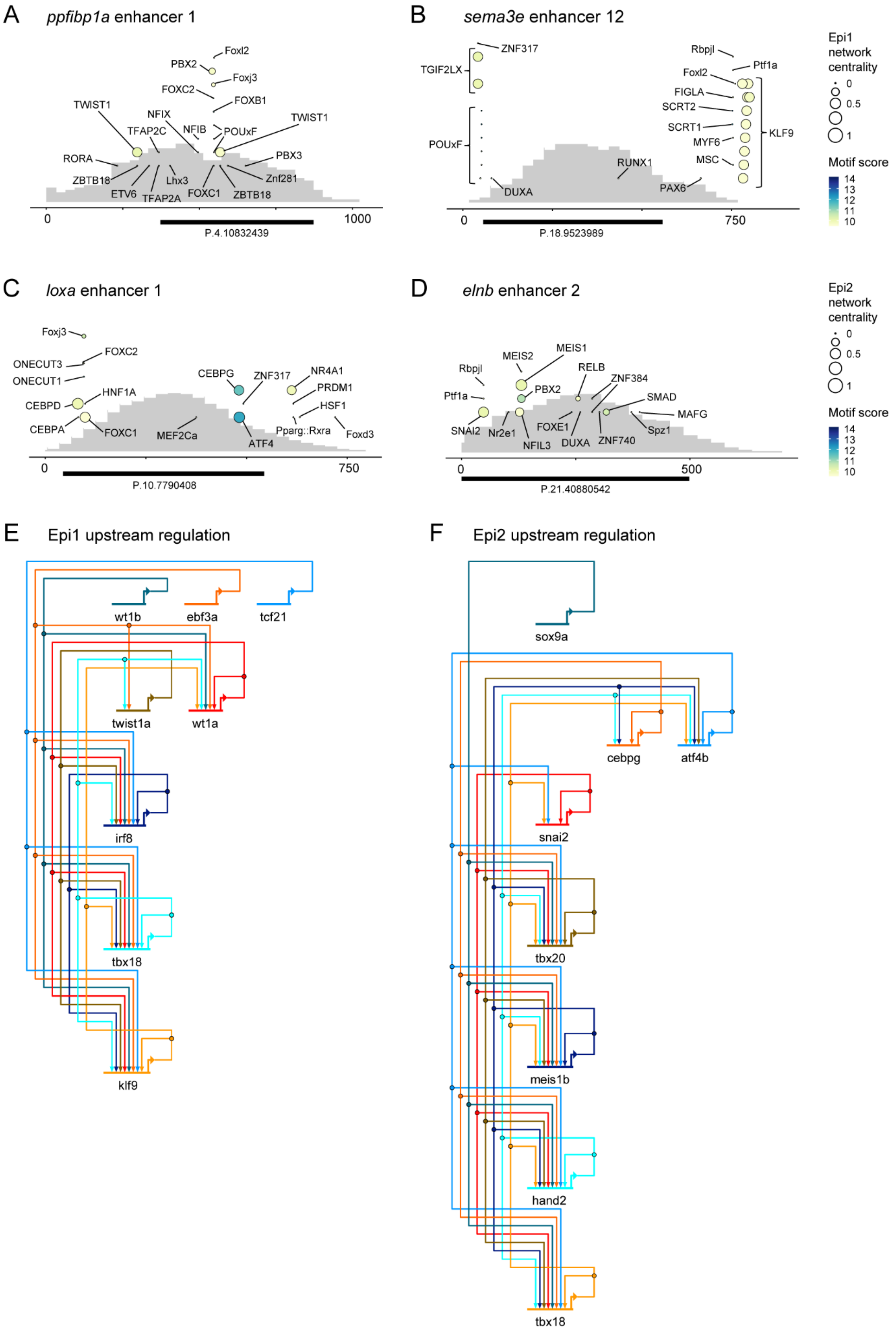
A
ppfibp1a enhancer 1
Foxl2
PBX2
Foxj3
FOXC2
FOXB1
TWIST1
NFIX
TWIST1
TFAP2C
NFIB
POUxF
PBX3
RORA
ZBTB18
Lhx3
Znf281
ETV6
TFAP2A
FOXC1
ZBTB18
0
1000
P.4.10832439
B
sema3e enhancer 12
ZNF317
Rbpjl
Ptf1a
TGIF2LX
Foxl2
FIGLA
SCRT2
SCRT1
KLF9
MYF6
POUxF
MSC
RUNX1
PAX6
DUXA
0
750
P.18.9523989
Epi1 network centrality
0
0.5
1
Motif score
14
13
12
11
10
C
loxa enhancer 1
Foxj3
FOXC2
ONECUT3
ONECUT1
CEBPG
ZNF317
NR4A1
PRDM1
HNF1A
CEBPD
HSF1
CEBPA
FOXC1
MEF2Ca
Pparg::Rxra
Foxd3
ATF4
0
750
P.10.7790408
D
elnb enhancer 2
MEIS1
MEIS2
Rbpjl
RELB
ZNF384
PBX2
Ptf1a
SMAD
MAFG
FOXE1
Spz1
SNAI2
Nr2e1
NFIL3
DUXA
ZNF740
0
500
P.21.40880542
Epi2 network centrality
0
0.5
1
Motif score
14
13
12
11
10
E
Epi1 upstream regulation
wt1b
ebf3a
tcf21
twist1a
wt1a
irf8
tbx18
klf9
F
Epi2 upstream regulation
sox9a
cebpg
atf4b
snai2
tbx20
meis1b
hand2
tbx18

**Figure S6. Transcriptomic regulation in larval epicardial subpopulations.**

(A) Transcription factor binding sites within the *ppfibp1a* enhancer 1 sequence. (B) Transcription factor binding sites within the *sema3e* enhancer 12 sequence. (C) Transcription factor binding sites within the *loxa* enhancer 1 sequence. (D) Transcription factor binding sites within the *elnb* enhancer 2 sequence. (E) Network schematic depicting predicted upstream transcriptional regulation between selected transcription factors in the Epi1 sub-network. (F) Network schematic depicting predicted upstream transcriptional regulation between selected transcription factors in the Epi2 sub-network. In A and B, bubble size indicates the eigenvector centrality of the factor in the tcf21 larval sub-network (In case of duplicated zebrafish genes, the centrality of the higher-scoring paralogue is indicated). Accessibility of the endogenous genomic region in the tcf21 larval condition is underlaid in grey. In C and D, bubble size indicates the centrality of the factor in the Epi2 sub-network. Accessibility of the endogenous genomic region in the tbx18 larval condition is underlaid in grey. In A-D, the quality of the binding sites is shown as colour-code. A black bar indicates the position of the endogenous peak region. Related to Figure 6.

| | Ensembl gene ID | Gene name | log2FC tcf21 larval vs tcf21 control | padj tcf21 larval vs tcf21 control | log2FC tcf21 cryo vs tcf21 sham | padj tcf21 cryo vs tcf21 sham | log2FC tcf21 larval vs tcf21 cryo | padj tcf21 larval vs tcf21 cryo |
|---|---|---|---|---|---|---|---|---|
| 1 | ENSDARG00000010752 | acsl4b | 1.741794 | 0.015754 | 3.928653 | 7.05E-07 | 0.887717 | 0.158975 |
| 2 | ENSDARG00000007709 | adamts8a | 3.312647 | 7.24E-06 | 3.221688 | 0.018312 | 0.024184 | 0.980812 |
| 3 | ENSDARG00000069027 | ADM_.2_of_2. | 3.780275 | 4.33E-09 | 7.063674 | 0.046071 | 7.577034 | 6.54E-19 |
| 4 | ENSDARG00000035552 | agtr2 | 4.624039 | 6.78E-11 | 7.592642 | 0.012542 | 4.407242 | 8.61E-11 |
| 5 | ENSDARG00000044056 | ahi1 | 1.671727 | 0.036223 | 1.652407 | 0.030173 | -1.60656 | 0.022271 |
| 6 | ENSDARG00000069463 | alox12 | 3.332651 | 0.000129 | 4.422566 | 0.022892 | 0.870902 | 0.246813 |
| 7 | ENSDARG00000025672 | antxr1a | 2.579563 | 0.011455 | 3.695372 | 0.018455 | -0.90985 | 0.308329 |
| 8 | ENSDARG00000077114 | arhgef16 | 2.827946 | 0.001043 | 7.476774 | 0.027759 | 0.929588 | 0.423479 |
| 9 | ENSDARG00000074335 | boc | 3.354693 | 4.21E-07 | 1.404823 | 0.035974 | 0.692769 | 0.288693 |
| 10 | ENSDARG00000088672 | CABZ01054962.1 | 4.008296 | 6.21E-06 | 5.982007 | 0.000836 | 0.561973 | 0.575888 |
| 11 | ENSDARG00000056477 | ccdc125 | 3.724221 | 1.26E-05 | 3.571746 | 0.024032 | 1.172339 | 0.197645 |
| 12 | ENSDARG00000078322 | col12a1a | 4.029554 | 1.85E-10 | 2.206892 | 0.00407 | -2.68553 | 0.000236 |
| 13 | ENSDARG00000099930 | col27a1b | 2.772146 | 0.000292 | 3.153061 | 5.27E-06 | -1.41577 | 0.050613 |
| 14 | ENSDARG00000075249 | fam171a2 | 2.974282 | 2.19E-05 | 2.089416 | 0.046971 | 0.548398 | 0.43068 |
| 15 | ENSDARG00000019815 | fn1a | 1.403613 | 0.04953 | 1.988468 | 0.000774 | -2.02383 | 0.001765 |
| 16 | ENSDARG00000002332 | frmd6 | 1.691401 | 0.036379 | 2.509336 | 0.022085 | -0.47873 | 0.530198 |
| 17 | ENSDARG00000027589 | fzd7b | 2.636029 | 0.000763 | 2.812548 | 0.012687 | 2.039187 | 0.001714 |
| 18 | ENSDARG00000068280 | GRB14 | 3.959977 | 1.70E-06 | 2.596079 | 0.038529 | 0.467341 | 0.513434 |
| 19 | ENSDARG00000040156 | grm4 | 2.176968 | 0.009614 | 5.753112 | 0.000399 | -0.6819 | 0.40064 |
| 20 | ENSDARG00000091650 | igflr1 | 1.67578 | 0.027484 | 9.425417 | 1.89E-05 | 0.191505 | 0.826787 |
| 21 | ENSDARG00000006314 | itgav | 1.741849 | 0.013289 | 5.314981 | 2.74E-07 | 0.582476 | 0.443503 |
| 22 | ENSDARG00000056998 | kirrelb | 3.556891 | 1.37E-05 | 3.863371 | 0.017198 | 0.072356 | 0.92437 |
| 23 | ENSDARG00000037960 | lrrc17 | 2.849457 | 0.000718 | 2.510102 | 0.023027 | 0.103044 | 0.920742 |
| 24 | ENSDARG00000098345 | mamdc2a | 2.686022 | 0.007423 | 7.664618 | 0.017014 | -0.12675 | 0.922047 |
| 25 | ENSDARG00000039034 | marcksl1a | 1.67189 | 0.02241 | 1.272369 | 0.027468 | -1.03347 | 0.128786 |
| 26 | ENSDARG00000036036 | mdka | 3.361961 | 7.33E-07 | 1.653534 | 0.024118 | -0.27189 | 0.749497 |
| 27 | ENSDARG00000017834 | micall2b | 2.235648 | 0.031424 | 5.631772 | 0.014674 | 0.146153 | 0.896862 |

| | | | | | | | | |
|---|---|---|---|---|---|---|---|---|
| 28 | ENSDARG00000078135 | MRC2 | 2.25452 | 0.019746 | 1.823538 | 0.015872 | -2.87414 | 3.04E-05 |
| 29 | ENSDARG00000053561 | ms4a17a.11 | 1.642945 | 0.033798 | 1.97186 | 0.028017 | 2.493893 | 0.000119 |
| 30 | ENSDARG00000038709 | MYADM_.1_of_2. | 4.557134 | 0.00019 | 3.701879 | 0.032333 | -0.44924 | 0.700834 |
| 31 | ENSDARG00000040700 | nab1b | 2.400334 | 0.009493 | 7.500825 | 0.018312 | 0.547463 | 0.628546 |
| 32 | ENSDARG00000005476 | nav3 | 1.900212 | 0.041068 | 1.453164 | 0.039781 | -3.76594 | 2.03E-07 |
| 33 | ENSDARG00000102855 | neo1a | 2.115014 | 0.005278 | 1.771404 | 0.038746 | -0.13104 | 0.870328 |
| 34 | ENSDARG00000014050 | ngfb | 3.339821 | 0.007657 | 2.699974 | 0.036565 | -2.28462 | 0.012833 |
| 35 | ENSDARG00000077474 | PLA2R1 | 2.373346 | 0.01224 | 3.755577 | 0.033168 | -0.50368 | 0.565361 |
| 36 | ENSDARG00000011821 | plod2 | 2.66647 | 0.000198 | 1.557711 | 0.034718 | 0.591285 | 0.424129 |
| 37 | ENSDARG00000059950 | plxdc2 | 1.575181 | 0.046839 | 1.700909 | 0.041726 | -1.60202 | 0.020833 |
| 38 | ENSDARG00000104267 | postnb | 3.632351 | 1.70E-08 | 1.823208 | 0.003158 | -2.62733 | 3.72E-05 |
| 39 | ENSDARG00000006604 | pvrl3b | 1.725769 | 0.036223 | 2.025319 | 0.037203 | -1.06411 | 0.131514 |
| 40 | ENSDARG00000036541 | rhbdf1a | 1.500691 | 0.036968 | 1.720341 | 0.013317 | -0.27069 | 0.685455 |
| 41 | ENSDARG00000004218 | rnd1l | 1.703733 | 0.012832 | 4.672439 | 0.03875 | 3.77165 | 8.64E-05 |
| 42 | ENSDARG00000033616 | sepn1 | 1.754733 | 0.025078 | 2.244371 | 0.002196 | -0.72414 | 0.335856 |
| 43 | ENSDARG00000098673 | si.ch211.233f10.6 | 4.110888 | 0.000779 | 2.686597 | 0.044178 | -1.87566 | 0.005085 |
| 44 | ENSDARG00000090901 | si.ch211.256e16.11 | 2.204378 | 0.015667 | 4.460419 | 0.046508 | -0.54781 | 1 |
| 45 | ENSDARG00000094327 | si.dkey.205k8.5 | 2.704398 | 0.018295 | 5.277347 | 0.038437 | -0.41637 | 0.687642 |
| 46 | ENSDARG00000097738 | si.dkey.31n5.4 | 3.354061 | 0.000369 | 7.085045 | 0.005685 | 1.009127 | 0.138889 |
| 47 | ENSDARG00000076899 | slc2a13b | 2.144187 | 0.046009 | 8.402399 | 0.002219 | -0.70363 | 0.543787 |
| 48 | ENSDARG00000103399 | SLC44A2_.2_of_2. | 2.66125 | 0.003083 | 6.148064 | 0.044844 | 1.478078 | 0.094355 |
| 49 | ENSDARG00000071304 | tbc1d2 | 2.25331 | 0.026858 | 3.657218 | 0.03661 | -0.76977 | 0.309954 |
| 50 | ENSDARG00000059363 | TGFBR2_.2_of_2. | 2.291925 | 0.004828 | 1.733537 | 0.016955 | -2.5614 | 0.000236 |
| 51 | ENSDARG00000045822 | tnnt2e | 2.916595 | 0.00533 | 8.023561 | 0.004158 | -0.42798 | 0.664572 |
| 52 | ENSDARG00000104082 | trps1 | 1.855579 | 0.03262 | 2.422432 | 0.036052 | -0.87757 | 0.221898 |

**Table S1.** Differential gene expression analysis results of genes enriched in larval epicardium (tcf21 larval) versus control, and in cryoinjured epicardium (tcf21 cryo) versus sham-treated epicardium (tcf21 sham). log2FC=log2 of fold change, padj=p-value adjusted for multiple comparisons.

| Enhancer name | GRCz10 coordinates | Primers |
|---|---|---|
| loxa_e1 | chr10:7790365-7791160 | forward: tcgagtttacgtaccgctagATCTTGAGAAGGTCCAGTTTCTTGAAC<br>reverse: TATCGCCGCAAGCTTgctagAATGGAAAACTGCAGCACTG |
| elnb_e2 | chr21:40880540-40881245 | forward: tcgagtttacgtaccgctagAAGTGGAGACCTAACATAACTCC<br>reverse: TATCGCCGCAAGCTTgctagATATATAAAGCTAAATCATATTCACTCTTGCAGC |
| ppfibp1a_e1 | chr4:10831835-10833341 | forward: CAACAGATCCCTCGActcgagCCCAGATGGCTTAAATTTCCTTC<br>reverse: tatcgccgcaAGCTTgctagcGCATGTGATAGCGTGGAGTG |
| sema3e_e12 | chr18:9523704-9524952 | forward: CAACAGATCCCTCGActcgagCACTCGCAATTCCTTACACAAT<br>reverse: tatcgccgcaAGCTTgctagcATGAGGGAAGAAGGGCAGAC |
| col12a1a_e1 | chr17:49653797-49655101 | forward: CAACAGATCCCTCGActcgagCTAAAGGAGTCAATGGCTGTTTT<br>reverse: tatcgccgcaAGCTTgctagcAACCAAGATGAAAGGCCGTG |
| mdka_e1 | chr7:38897699-38898662 | forward: CAACAGATCCCTCGActcgagTGGGTGTCAGGCTATTGTTT<br>reverse: tatcgccgcaAGCTTgctagcACAACACAAAGCTCTGACCC |

**Table S2.** Primers used to amplify putative enhancer regions in the GRCz10 genome assembly. Underlined sequences are homology regions used to insert the enhancer sequences into the Ac/Ds reporter vector.
